# Pipefish locally adapted to low salinity in the Baltic Sea retain phenotypic plasticity to cope with ancestral salinity levels

**DOI:** 10.1101/2020.11.12.379305

**Authors:** Henry Goehlich, Linda Sartoris, Kim Sara Wagner, Carolin C. Wendling, Olivia Roth

## Abstract

Genetic adaptation and phenotypic plasticity facilitate the invasion of new habitats and enable organisms to cope with a rapidly changing environment. In contrast to genetic adaptation that spans multiple generations as an evolutionary process, phenotypic plasticity allows acclimation within the life-time of an organism. Genetic adaptation and phenotypic plasticity are usually studied in isolation, however, only by including their interactive impact, we can understand acclimation and adaptation in nature. We aimed to explore the contribution of adaptation and plasticity in coping with an abiotic (salinity) and a biotic (Vibrio bacteria) stressor using six different populations of the broad-nosed pipefish Syngnathus typhle that originated from either high or low saline environments. We hypothesized that wild S. typhle populations are locally adapted to the salinity and prevailing pathogens of their native environment, and that short-term acclimation of parents to a novel salinity may aid in buffering offspring phenotypes in a matching environment. To test these hypotheses, we exposed all wild caught animals, to either high or low salinity, representing native and novel salinity conditions and allowed animals to mate. After male pregnancy, offspring was split and each half was exposed to one of the two salinities and infected with Vibrio alginolyticus bacteria that were evolved at either of the two salinities in a fully reciprocal design. We investigated life history traits of fathers (offspring survival, offspring size) and expression of 47 target genes in mothers and offspring.

Pregnant males originating from high salinity exposed to low salinity were highly susceptible to opportunistic fungi infections resulting in decreased offspring size and number. In contrast, no signs of fungal infection were identified in fathers originating from low saline conditions suggesting that genetic adaptation has the potential to overcome the challenging conditions of low salinity. Genetic adaptation increased survival rates of juveniles from parents in lower salinity (in contrast to those from high salinity). Juvenile gene expression indicated patterns of local adaptation, trans-generational plasticity and developmental plasticity. The results of our study suggest that pipefish locally adapted to low salinity retain phenotypic plasticity, which allows them to also cope with ancestral salinity levels and prevailing pathogens.

## 1. Introduction

Genetic adaptation and phenotypic plasticity (Chevin, Lande et al. 2010) facilitate the invasion of new habitats and permit coping with the consequences of global climate change (Brierley and Kingsford 2009, Poloczanska, Brown et al. 2013, Urban 2015). Genetic adaptation is a multigenerational process spreading in the population over the rise and fixation of novel mutations (Chatterjee, Pavlogiannis et al. 2014), or over selection on standing genetic variation and shifts in allele frequency (Barrett and Schluter 2008, Eizaguirre, Lenz et al. 2012, Torda, Donelson et al. 2017). In contrast, phenotypic plasticity is an individual trait that enables organisms of one genotype to show multiple, alternative phenotypes in response to biotic or abiotic conditions (West-Eberhard 1989). The environment influences the phenotype (Chevin, Lande et al. 2010) and elicits changes in gene expression, which can then impact individual development, morphology, physiology and behavior (Angers, Castonguay et al. 2010). Phenotypic responses can occur within the life-time of an organism (reversible and developmental plasticity) or persist across one or several generations (trans-generational plasticity).

Trans-generational plasticity (TGP) is the non-genetic inheritance of an alternative phenotype by transferring nutrients, hormones, proteins or epigenetic marks from the parent to the offspring generation (Sunday, Calosi et al. 2014). The impact of TGP may differ among species, life stages and abiotic conditions (Uller, Nakagawa et al. 2013, Laland, Uller et al. 2014) as well as the biotic interaction partners (e.g. parasite type or strain) (Beemelmanns and Roth 2016, Beemelmanns and Roth 2016, Beemelmanns and Roth 2017, Roth, Beemelmanns et al. 2018). TGP can be adaptive and result in increased offspring performance when environmental conditions of parental and offspring generations match (Sunday, Calosi et al. 2014). This has been shown for instance in wild Atlantic silversides exposed to ocean acidification (Murray, Malvezzi et al. 2014) or in three-spine sticklebacks exposed to heat stress (Shama and Wegner 2014). However, TGP can also induce negative carry-over effects (Eriksen, Bakken et al. 2006, Marshall 2008), e.g. increased mortality in the early life stages of sticklebacks upon changes in salinity levels (Heckwolf, Meyer et al. 2018). Adaptive phenotypic plasticity allows organisms to survive and reproduce in a novel environment but was suggested to slow down genetic adaptation by buffering against the effects of natural selection (Kelly 2019).Whether TGP is enhancing or constraining adaptation is still debated and may depend on various factors such as species, traits or the level of current environmental variability and predictability (Reed, Waples et al. 2010, Lind, Zwoinska et al. 2020).

The interactive contribution of genetic adaptation and phenotypic plasticity in invading new habitats and coping with climate change has been rarely addressed, instead, the two mechanisms were mainly studied in isolation (Gienapp, Teplitsky et al. 2008). However, to depict and understand biological responses to climate driven environmental changes, we need models (Donelson, Sunday et al. 2019) and experiments (Kelly 2019) addressing such mechanisms simultaneously. An approach to study the interaction between genetic adaptations and phenotypic plasticity are space-for-time experiments (Blois, Williams et al. 2013, Kelly 2019), where organisms living along a natural gradient can serve as a prediction for how organisms can cope with future environmental conditions (Reusch, Dierking et al. 2018).

Even though salinity shifts are predicted to have strong implications for coastal populations (Meier, Kjellstrom et al. 2006, Andersson, Meier et al. 2015, Kniebusch, Meier et al. 2019), the main focus of climate change research still lies on warming and acidification studies (but see DeFaveri and Merila 2014, Hasan, DeFaveri et al. 2017, Heckwolf, Meyer et al. 2018). Changing ocean salinity will have major impacts on coastal and polar ecosystems (Gibson and Najjar 2000, Loder, van der Baaren et al. 2015), because of the overriding effects on the physiology of aquatic organisms (Morgan and Iwama 1991, Velasco, Gutierrez-Canovas et al. 2019), comprising metabolism, growth, development, immunity and reproduction in teleost fishes (Haddy and Pankhurst 2000, Boeuf and Payan 2001). Osmoregulation enables marine organisms to acclimate to different salinity levels but consumes up to 50% of the fish’s total energy budget (Boeuf and Payan 2001). High energy demand for osmoregulation results in metabolic trade-offs (DeWitt, Sih et al. 1998), which makes genetic adaptation to novel salinities important.

The Baltic Sea is particularly prone to future reductions in salinity due to little water exchange with the North Sea and river runoffs from the surrounding countries. Increased precipitation in the northern part may cause a decrease by up to 30% in surface salinity by the end of the century (Meier, Kjellstrom et al. 2006, Andersson, Meier et al. 2015). Already today, the Baltic Sea is characterized by a strong salinity gradient ranging from 30 PSU in the transition to the North Sea to an almost freshwater environment in the north-eastern parts making it an ideal setting for space for time experiments (Blanquart and Gandon 2013, Heckwolf, Meyer et al. 2018). The stability of the salinity gradient (Janssen, Schrum et al. 1999, Hinrichs, Jahnke-Bornemann et al. 2019), the energetic cost of both, osmoregulation (Boeuf and Payan 2001) and phenotypic plasticity (DeWitt, Sih et al. 1998), promote genetic adaptation in teleost fishes towards different salinity levels in the Baltic Sea (DeFaveri and Merila 2014, Berg, Jentoft et al. 2015, Guo, DeFaveri et al. 2015, Guo, Li et al. 2016). If salinity levels in the new environment are relatively stable, genetic assimilation was suggested to result in reduced plasticity and more adaptive genotypes (Angers, Castonguay et al. 2010). Adaptation to the low salinity conditions of the Baltic Sea and the isolation from the Atlantic source population is also accompanied by a loss of genetic diversity (Johannesson and Andre 2006, Holmborn, Goetze et al. 2011). Therefore, adaptation to low salinity may result in reduced osmoregulatory plasticity, such as changes in kidney morphology and gene expression (Hasan, DeFaveri et al. 2017), and thus hamper the ability to cope with further salinity fluctuations. TGP may not be sufficient to buffer the negative impacts of salinity change (Heckwolf, Meyer et al. 2018), in particular if salinity is subject to strong fluctuations and if populations are locally adapted. However, increased selection due to negative carry-over effects may facilitate rapid adaptation but may also reduce genetic variation, which raises the risk of extinction (Heckwolf, Meyer et al. 2018). Unclear remains whether strong selection for genetic adaptation to low saline environments generally resulted in a reduction of phenotypic plasticity or whether, alternatively, animals have evolved different strategies to cope with salinity changes.

A suitable organism to study the interactive contribution of genetic adaptation and phenotypic plasticity is the sex-role reversed broad-nosed pipefish Syngnathus typhle (Syngnathidae, Teleostei). S. typhle inhabits a wide range of waters with different salinity levels along the European coastline from the Black Sea in Eastern Europe to the Mediterranean Sea and from the Eastern Atlantic to the north of the Baltic Sea (Wilson and Veraguth 2010). TGP in response to immune and temperature challenges has been demonstrated in broad-nosed pipefish in numerous studies (Beemelmanns and Roth 2016, Beemelmanns and Roth 2017, Roth and Landis 2017) as well as the impairing effect of low salinity on the immune system (Birrer, Reusch et al. 2012). Beyond the direct impact of salinity changes on organisms and populations (genotype x environment interaction, GxE), salinity shifts may increase or decrease the virulence of parasites and pathogens (genotype x genotype x environment interaction, GxGxE) (Stockwell, Purcell et al. 2011, Hall, Vettiger et al. 2013, Poirier, Listmann et al. 2017) and alter co-evolutionary dynamics between host and pathogens (Mostowy and Engelstadter 2011, Molnar, Kutz et al. 2013, Brunner and Eizaguirre 2016, Kutzer and Armitage 2016).

The abundance and virulence of opportunistic and omnipresent marine pathogens, such as several strains of the Vibrio bacteria clade (Baker-Austin, Trinanes et al. 2017) are modulated by salinity and temperature (Chen, Li et al. 2011, Oberbeckmann, Wichels et al. 2011, Baker-Austin, Trinanes et al. 2017). Vibrio alginolyticus frequently infects pipefish in the Baltic Sea (Roth, Keller et al. 2012) and is known to cause higher mortality in artemia and herring at low salinity (Dayma, Raval et al. 2015, Poirier, Listmann et al. 2017). Increases in bacterial virulence are evoked due to a combination of phenotypic changes, including bacterial biofilm formation (Dayma, Raval et al. 2015, Kim and Chong 2017) and the expression of bacterial motility and virulence factors (Hase and Barquera 2001). We hypothesized that genetic adaptation of the pipefish to local salinity and the prevailing pathogens may compensate for the previously observed drop of immunological activity in case of exposure to decreasing salinities (Birrer, Reusch et al. 2012, Poirier, Listmann et al. 2017) and, hence, has the potential to reduce the negative impact of pathogens like Vibrio bacteria (Roth, Keller et al. 2012).

To explore how pipefishes have genetically adapted to long-term salinity changes and how this adaptation influences their phenotypic plasticity to cope with short-term shifts in salinity, we compared the potential of pipefish originating from either high or low salinity environments to react towards salinity shifts with developmental and trans-generational plasticity. Furthermore, we investigated how adaptation and acclimation of the pipefish host and the bacterial Vibrio pathogen to high and low salinity changes the host-pathogen interaction. We tested the following hypotheses: 1) S. typhle populations are genetically adapted to the salinity in their local habitat, 2) adaptive trans-generational plasticity in matching parental and offspring salinity results in enhanced juvenile survival and matching gene expression pattern in the parental and offspring generation, 3) S. typhle populations locally adapted to low salinity have reduced phenotypic plasticity and are not able to cope with ancestral salinity levels, and 4) bacterial virulence is higher at low salinity.

To investigate how S. typhle have adapted towards their local salinity and local pathogens in the past (genetic adaptation) and to assess their consecutive acclimation potential (phenotypic plasticity) towards salinity shifts and their immune response towards a bacterial infection, we collected six S. typhle populations in the Baltic Sea. Fish were collected at three sampling sites with high saline conditions and at three sampling sites with low saline conditions. In a laboratory aquaria experiment, animals were exposed to either their native salinity (high or low respectively) or the salinity of the other three populations (novel salinity). Upon successful male pregnancy, offspring were exposed to either native or novel salinity conditions, in a fully reciprocal design. Subsequently, juvenile fish were injected with a V. alginolyticus strain that evolved for 90 days either at low or high salinity in the laboratory. In addition to life history traits and mortality, we investigated the expression of 47 target genes involved in (i) general metabolism, (ii) immune response, (iii) gene regulation (DNA and histone modification) and (iv) osmoregulation.

## 2. Material and Methods

### 2.1 Sampling of adult pipefish populations

The parental Syngnathus typhle generation was caught in seagrass meadows of six sampling sites along the German coastline of the Baltic Sea in spring 2017 (Figure 1 & Table 1). Three sampling sites are characterized by relatively high salinity conditions (14 - 17 PSU; high salinity origin; H) and three sampling sites by relatively low salinity conditions (7 - 11 PSU; low salinity origin, L; Table 1). Salzhaff was assigned the category low because salinity drops are common after rainfall accompanied with freshwater discharge due to enclosed morphology of the inlet. Therefore, pipefish in Salzhaff are likely to be exposed to salinity levels below 10 PSU. A minimum of 30 non-pregnant males and 30 females were caught snorkeling with hand nets at each sampling site at depths ranging between 0.5 and 2.5 m. At each sampling site, water temperature and salinity were measured from water collected about 1 m below the surface using a salinometer (WTW Cond 330i).

**Figure 1:**
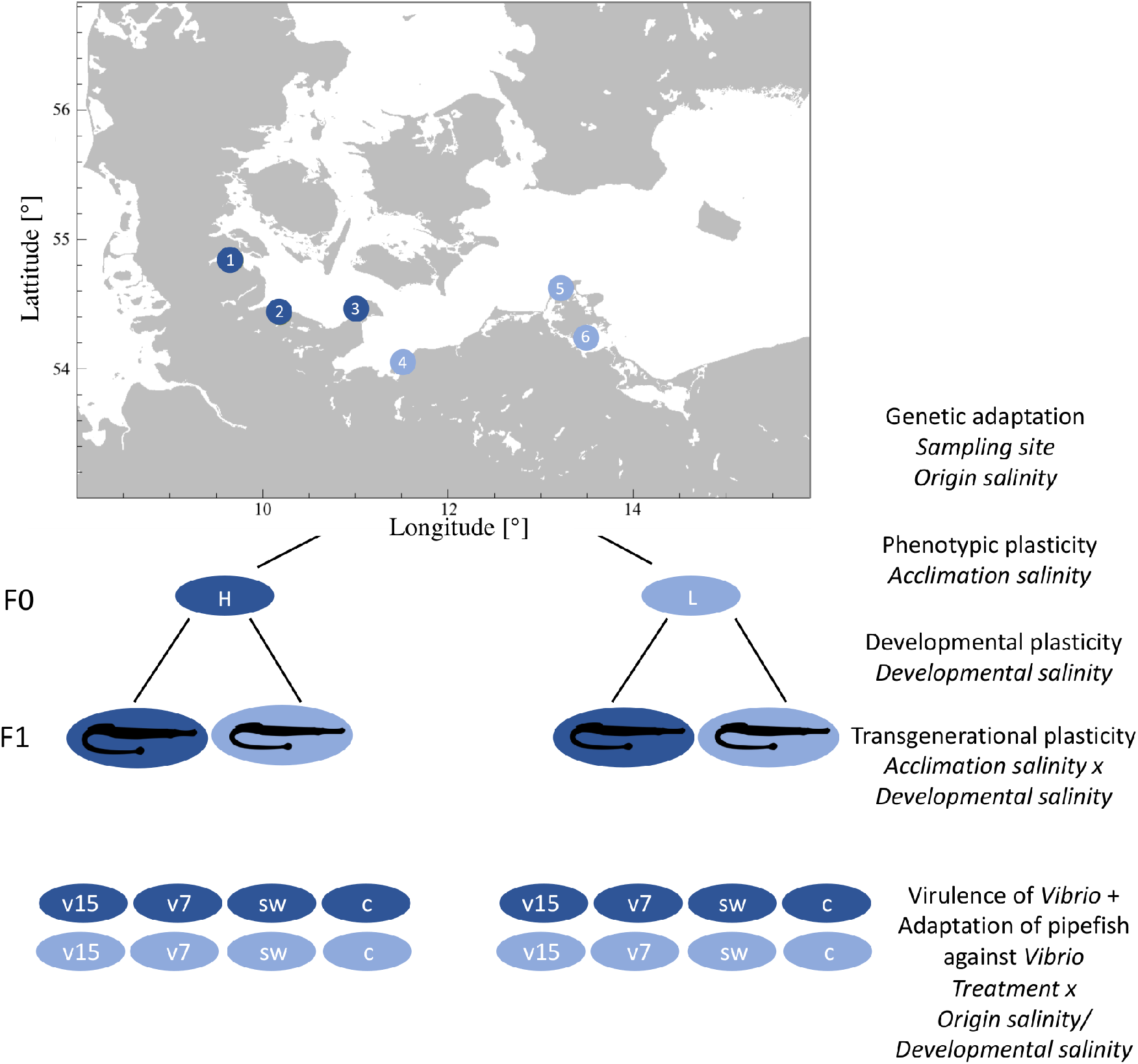
Fully reciprocal experimental design. We sampled pipefish along the Baltic Sea coast, at three sampling sites from a relatively high saline environment (high origin salinity: 14 - 17 PSU; dark blue circles; subsequently labeled as italic H): 1) Flensburg Fjord, 2) Falckenstein Strand and 3) Fehmarn and three sampling sites with a relatively low salinity level (low origin salinity: 7 - 11 PSU; light blue circles; subsequently labeled as italic L): 4) the Salzhaff and 5) Ruegen North and 6) Ruegen South. In the laboratory males and females were kept separately and acclimated to the opposing salinity (acclimation salinity: 15 PSU (H, dark blue), 7 PSU (L, light blue)) or their respective native salinity. Subsequently, males and females were allowed to mate and pregnant males were kept at constant conditions. Half of the F1 generation was either exposed to high (h) or low (l) salinity within 24h after birth (developmental salinity). Ten days post hatch juveniles were injected with Vibrio alginolyticus evolved at 15 PSU (v15) or at 7 PSU (v7), sham injected with sterile seawater (W) or left naive (C) (treatment). Label in italic on the right side correspond to the factors that were considered in the statistical models.

**Table 1:** Pipefish sampling sites with coordinates, sampling date and ambient salinity and water temperature.

| **Sampling sites**<br>**(Abbreviation)** | **GPS Coordinates** | **Salinity**<br>**(PSU)** | **Salinity**<br>**(Category)** | **Water**<br>**Temperature**<br>**(°C)** |
| --- | --- | --- | --- | --- |
| Flensburg Fjord<br>Westerholz (Flens) | 54°49'14 N<br>9°40'26 E | 17 | High | 15 |
| Kiel Fjord<br>Falckensteiner Strand (Falck) | 54°49'26 N<br>10° 11'30 E | 14 | High | 10-11 |
| Orth Bay<br>Fehmarn (Fehm) | 54°26'55 N<br>11° 3'19 E | 15 | High | 13 |
| Salzhaff<br>Werder (Salz) | 54° 1'35 N<br>11° 3'57 E | 10-11 | Low | 14 |
| Wieker Bodden<br>Wiek (RuegN) | 54° 37'20 N<br>13° 16'56 E | 8 | Low | 18 |
| Strelasund<br>Grabow (RuegS) | 54° 13'32 N<br>13° 24'25 E | 7 | Low | 12 |

Pipefish were transported in large aerated coolers to the aquaria facilities of the GEOMAR (Westshore) in Kiel (Germany). Females and males were separated and kept in groups of 5-7 per 80-liter tank resulting in a total number of 36 tanks. These tanks were connected to two independent circulating water systems containing either high saline (15 PSU; n = 18) or low saline water (7 PSU, n = 18) and equipped with artificial seagrass. Pipefish from high salinity origins were kept at 15 PSU (Baltic Sea water) and those from low salinity origins at 7 PSU (Baltic Sea water, diluted with deionized water and tap water (ratio 2:1:1) to keep the water alkalinity constant). The water temperature throughout the experiment was 18° C and illumination was set to a 16:8 h day and night cycle. Pipefish adults were fed twice a day with frozen and occasionally with live mysids.

After pipefish were acclimated to laboratory conditions for at least two days, half of the individuals from each sampling site were gradually acclimated to the novel salinity over four days. Each day, tanks were briefly connected to the 15 PSU or the 7 PSU circulating system to either increase or decrease the salinity by 1.5 to 2 PSU. The other half of the fish remained in their native salinity. Apart from the salinity adjustment, all 36 tanks remained disconnected from the circulation system during the time of salinity acclimation. Subsequently, four to six randomly chosen males and four to six females originating from the same sampling site and acclimated to the same salinity, were placed together in one of the 36 tanks connected to circulating water systems of either high or low acclimation salinity (Figure 1). During mating and male pregnancy, fish maintenance and aquaria set-up remained as previously described.

One week after mating, pipefish males started to show signs of infection with a fungus growing inside and on the brood pouch. Three weeks after mating, we visually assessed and photographically documented the prevalence of fungus.

### 2.2 Sampling of adult pipefish for targeted gene expression & population genetics

Four days after mating, females were removed from the tanks and immediately euthanized using anesthetic tricaine methane sulfonate (MS-222, 500 mg/L). We measured standard body length and total weight and removed the gills to store them in RNAlater at 4° C overnight and subsequently at −80° C. Fin clips were taken and placed in 96% Ethanol for population genetic analysis.

### 2.3 Population genetics using microsatellites

#### 2.3.1 DNA isolation & preparation

Genomic DNA was isolated from fin clips of F0 female pipefish using the DNeasy 96 Blood & Tissue Kit (Qiagen, Venlo, Netherlands) following the manufacturer’s protocol. All samples were incubated and eluted twice to obtain a higher extraction yield. A subset of the isolated genomic DNA was quantified using NanoDrop (Spectrometer; Peqlab, Erlangen, Germany) and visually evaluated by gel electrophoresis on a 1.2% agarose gel (GelRed nucleic acid stain, Lambda DNA/HindIII Marker and 1kb DNA marker (Invitrogen; Thermo Fisher Scientific, Germany)).

All 144 *S. typhle* samples were genotyped for 11 microsatellite loci, with a minimum of 20 individuals per sampling site. Genotyping was performed in three pooled reactions, each containing 3-4 primer pairs that were designed on an expressed sequence tag (EST) library of *S. typhle* (**Pool A**: Sy_ty_1, Sy_ty_4, Sy_ty_6, Sy_ty_7; **Pool B**: Sy_ty_11, Sy_ty_22, Sy_ty_23; **Pool C**: Sy_ty_16, Sy_ty_17, Sy_ty_21, Sy_ty_24 (Jones, Rosenqvist et al. 1999, Roth, Keller et al. 2012)). Microsatellites and the associated primer pairs and the Multiplex PCR protocol can be found in GenBank under accession numbers JQ598279– JQ598290. Primers had an initial concentration of 5 pmol and were color labeled with either Hex green or Fam blue to allow differentiation during fragment analysis. In a 10 *μ*l reaction, several loci were amplified simultaneously from 1 *μ*l of extracted DNA using 5 *μ*l of the Multiplex PCR Master Mix (Qiagen) and varying amounts of the pooled primer mixes (Pool A: 1.75 *μ*l, Pool B: 0.75 *μ*l, Pool C: 1.5 *μ*l). Three negative controls (ddH_2_O) were added onto each 96-well plate.

Capillary electrophoresis and fragment analysis were performed using the 3130xl Genetic Analyzer (Applied Biosystems/Thermo Fisher Scientific). A loading mix containing 8.75 *μ*l HiDi Formamide and 0.25 *μ*l GeneScan 350 ROX dye Size Standard (Applied Biosystems/Thermo Fisher Scientific) was added to 1 *μ*l of each PCR product. Prior to the fragment run, samples were denatured in a thermo cycler for 2 min at 90° C.

#### 2.3.2 Microsatellite analysis

Raw fragment data were scored using the GeneMarker Genotyping Software (Liu et al. 2011). The software displays allele frequency panels that identify the alleles for each locus in each sample, thus provides an overview of whether individuals are homozygous or heterozygous for certain alleles at a locus. Additionally, the raw data were screened using the Microsatellite Data Checking Software Micro-Checker (Van Oosterhout et al. 2004). Micro-Checker identifies genotyping errors caused by non-amplified null-alleles that either appear due to mutations in the primer binding regions or generally occur in fragment analysis because PCR shows greater efficiency in longer sequences. GENETIX (Belkhir et al. 2004) was used to describe the level to which the genotype frequency differed from the expected Hardy-Weinberg equilibrium (HWE) frequency by calculating a global FST value as a correlation of inbreeding in the substructure vs. in the entire population. For completeness, pairwise FST values were calculated to display distances between pairs of haplotypes and a FIS value was calculated as a correlation of inbreeding vs. random mating within the population. Although GENETIX has a greater statistical power, the population structure within the multi-locus genotype data was further investigated by the STRUCTURE Software for Population Genetics Inference (Pritchard et al. 2000). Based on the Bayesian clustering method, STRUCTURE creates an admixture model, which provides likelihood scores for each individual of belonging to a certain population. The model was tested with varying numbers of expected populations ranging from a minimum of two (high salinity vs. low salinity) to a maximum of six (number of sampling stations). Visualization of the population clustering was performed using the PHYLogeny Inference Package PHYLIP (Felsenstein 1989). PHYLIP provides a pipeline of programs to randomize comparisons, create randomized trees, which are then assembled to a final phylogeographic tree that is based on the most frequent combinations found within the randomized trees. As the retrieved fragment data did not provide any lineage data that allows to draw conclusions with regard to a common ancestor, we created an unrooted phylogeographic tree.

### 2.4 Candidate gene expression of females

To assess local adaptation to salinity and the potential of S. typhle to cope with novel salinity conditions, we selected candidate genes from three different functional categories, i.e. (i) immune response, (ii) metabolism and (iii) gene regulation (DNA and histone modification) (Supplement 1 (Table S1)). Immune genes were further subdivided into innate, adaptive and complement system genes and gene regulation genes into activating and silencing genes.

#### 2.4.1 RNA extraction and reverse transcription

RNA was extracted from gill tissue of adult pipefish that was stabilized in RNAlater using the RNeasy® Universal Tissue kit (QIAGEN, Venlo, Netherlands). Tissue samples were homogenized by adding a 5 mm stainless steel bead into each collection tube and placing them into a homogenizer shaking for two times 30 seconds at 25 Hertz. Thereafter, we followed the manufacturer’s protocol “Purification of Total RNA from Animal Tissues Using Spin Technology”. RNA concentration (extraction yield) and purity of the samples were checked by spectrophotometry (NanoDrop ND-1000 Spectrometer; Peqlab, Erlangen, Germany). Protein contamination was quantified using the adsorption ratio of 260/280 nm (target > 2.0) and the ratio 260/230 nm (target > 1.8) was used to detect organic contamination. A fixed amount of RNA (300 ng/sample = 50 ng/*μ*l) was then reverse transcribed into cDNA using the QuantiTect Reverse Transcriptase kit (QIAGEN, Venlo, Netherlands).

#### 2.4.2 Preamplification of cDNA and candidate gene expression

For each sample, 1.4 *μ*l target cDNA was pre-amplified with 0.5 *μ*l primer pool mix of all 48 genes (500 nM), 2.5 *μ*l TaqMan PreAmp Master Mix (Applied Biosystems, Waltham, MA, USA) and 0.7 *μ*l H2O (10 min at 95° C, 14 cycles: 15 sec at 95° C followed by 4 min at 60° C). Afterwards, the PCR product was diluted 1:10 with low TE buffer (10 mM Tris, 0.1 mM EDTA, pH 8). The sample mix for the 96.96 Dynamic ArrayTM IFCs chips contained 3.1 *μ*l pre-amplified and diluted PCR product, 3.55 *μ*l SsoFast-EvaGreen Supermix with Low ROX (Bio-Rad Laboratories, Hercules, CA, USA) and 0.37 *μ*l 20 x DNA Binding Dye Sample Loading Reagent (Fluidigm) per sample. The assay mix for the chip contained 0.7 *μ*l primer pair mix (50 *μ*M), 3.5 *μ*l Assay Loading Reagent (Fluidigm) and 2.8 *μ*l low TE buffer per primer pair. Chips were loaded with 5 *μ*l sample mix and 5 *μ*l assay mix. To measure gene expression, the chips were placed into the BioMark system (Fluidigm, South San Francisco, CA, 15 USA) applying the ‘GE fast 96×96 PCR+Melt v2.pcl’ protocol (Fluidigm). Each of the chips contained two technical replicates per sample and gene, two no-template controls (H2O), one control for gDNA contamination (-RT) and one between plate control.

### 2.5 Juvenile infection experiment

#### 2.5.1 Experimental design and treatment groups

Within the first 24 hours after birth, half of the juveniles from each clutch was exposed to native salinity conditions and half to novel salinity conditions in a fully reciprocal design. Juveniles were fed twice a day with freshly hatched, nutrient enriched (Aqua Biotica orange+TM) Artemia salina nauplii. Siblings were kept together in one non-aerated 1.5 l tank, of which one third of the water was exchanged daily. Once a day, left-over food was removed using single-use pipettes and mortality was documented.

Ten days post-hatch, juveniles received one of the four following treatments: i) no injection (c), ii) sham injection of autoclaved seawater with the equivalent salinity, i.e. 15 or 7 PSU (sw) or iii) injection of Vibrio alginolyticus strain K01M1, which evolved for 90 days under laboratory condition either at 15 (v15) or 7 PSU (v7) (Goehlich, Roth et al., unpublished data). 2 *μ*l of sterile seawater with or without bacteria was injected in the ventral part of the juveniles, using a MonojectTM insulin syringe (Coviden) with a sterile 30 Gauge needle. Subsequently, all juvenile siblings with the same treatment were placed in one 500 ml Kautex bottle containing seawater with the respective salinity of the 1.5 l tanks. Survival of juveniles was documented for six days and fish maintenance was according to the procedure described for 1.5 l tanks. One day post infection, one juvenile from each treatment (Kautex bottle) was euthanized and decapitated to assess expression of candidate genes. Standard body length was measured and whole-body samples were stored in RNAlater overnight at 4° C and subsequently at −80° C.

#### 2.5.2 Characterization and evolution of *Vibrio alginolyticus* strain used for injection

The Vibrio alginolyticus strain K01M1 used for injection of pipefish juveniles was isolated from a healthy pipefish caught in the Kiel Fjord (Roth, Keller et al. 2012) and fully sequenced (Chibani, Roth et al. 2020). The strain was evolved for 90 days either at 15 or 7 PSU (medium 101: 0.5% (w/v) peptone, 0.3% (w/v) meat extract, 1.5% (w/v) or 0.7% (w/v) NaCl in Milli-Q deionized water) (Goehlich, Roth et al., unpublished data). We used the same strain and evolved it at two different salinities to ensure that salinity is the only driver for differences in bacterial virulence, which can potentially also be influence by the presence of filamentous phages (Waldor and Mekalanos 1996, Ilyina 2015, Chibani, Hertel et al. 2019).

After 90 days the bacterial populations were diluted and plated onto Vibrio selective Thiosulfate-Citrate-Bile-Saccharose (TCBS) agar plates (Fluka AnalyticalTM). The next day, single colonies from each plate were picked and grown overnight in medium 101 with the respective salinity. Subsequently, cultured bacteria were stored at −80°C as 33% glycerol stocks. For the infection experiment, part of the glycerol stocks were plated onto TCBS agar and one clone was grown in a 50 ml Falcon tube containing 30 ml medium 101 in the respective salinity for 24 hours, at 25 °C with shaking at 230 rpm. Overnight cultures were centrifuged for 20 min at 2000 rpm. The supernatant was discarded and the cell pellet was resuspended in 3 ml sterile seawater (7 or 15 PSU respectively) to achieve similar bacterial densities of 5*1010 ml-1.

#### 2.5.3 Juvenile gene expression

We measured gene expression of juveniles to assess the effect of (a) genetic adaptation (i.e. origin salinity) on gene expression, (b) trans-generational effects driven by an interaction between acclimation salinity and developmental salinity and (c) developmental plasticity induced by developmental salinity. Furthermore, we investigated (d) whether virulence differed in V. alginolyticus evolved at 15 or 7 PSU and whether juveniles from parents originating from a matching salinity were better adapted to Vibrio strains evolved at the respective salinity. Therefore, we selected genes from three functional categories, namely (i) immune response (ii), general metabolism (iii) gene regulation (DNA and histone modification) as described above for female pipefish S. typhle (Section 2.4). Compared to female gene expression, eleven genes from the categories (i)-(iii) were replaced by osmoregulation genes (iv). We selected osmoregulatory genes from teleost studies (S 2) and designed specific primers with Primer3Web (Koressaar and Remm 2007, Untergasser, Cutcutache et al. 2012) (S 3). RNA extraction and quantification of gene expression were conducted as described before with the following modifications due to a higher RNA yield: the fixed amount of RNA that was reverse transcribed into cDNA was 400 ng/sample (67 ng/*μ*l) instead of 300 ng/sample (50 ng/*μ*l) and pre-amplified cDNA was diluted 1:10 and instead of 1:20.

### 2.6 Statistics

All statistical analyses and visualizations were performed in the R 3.6.1 environment (RCoreTeam 2020).

#### 2.6.1 Life history traits

We used two-way ANOVAs to assess size and weight differences between adults as well as differences in clutch size and in total length between juveniles at 10 days post-hatch. Fixed factors included *origin salinity* (Salinity at sampling sites of origin two levels: *High* or *Low*), *acclimation salinity* (High or Low), *sex* of the pipefish (male or female) and the *sampling site* (Flens, Fehm, Falck, Salz, RuegN or RuegS) nested in *origin salinity*. ANOVA of clutch size additionally included the average body length of males exposed to a given treatment. Homogeneity of variances was tested by Fligner test and normal distribution of data by using the Cramer-von Mises normality test. The clutch size was square root transformed to achieve normal distribution of residuals.

We performed two spearman-rank correlations using the function ggscatter (package: ggpubr) to test for 1) correlation between the total length of adult pipefish and the salinity measured at the sampling site on the day of capture as well as 2) between clutch size and average male size of sampling site. The clutch size of males originating from high salinity and acclimated to low salinity conditions was removed from the correlation due to fungus infection. Post-hoc tests were carried out using Tukey’s “honest significant difference” (TukeyHSD, package: multcomp) (Hothorn, Bretz et al. 2020). Gene expression of parental generation & juveniles

#### 2.6.2 Gene expression of parental generation & juveniles

From the Fluidigm output data, mean cycle time (Ct) and standard deviation (SD) for each of the two technical replicates were calculated. Expression measurements with a coefficient of variation (CV; CV = SD/Ct) larger than 4% were excluded from the study (Bookout and Mangelsdorf 2003). For females, the combination of HDAC1 and HDAC3 were identified as the optimal reference genes (geNorm V < 0.15 (Vandesompele, De Preter et al. 2002)) with high reference target stability (geNorm M ≤ 0.5), based on 155 samples (S 5) and 34 target genes in *Qbase*+*3.0* (Biogazelle; Hellemans, Mortier et al. 2007).

In the analyses of juvenile gene expression one osmoregulation gene (15% NAs) and 36 samples were excluded from the study due to failed reactions on the Fluidigm chip in at least one of the duplicates. Samples with more than 10% excluded genes were omitted from the analysis, as many missing Ct values are indicative for insufficient sample quality. Remaining missing Ct values were substituted by the mean Ct for the given gene calculated from all other samples, as subsequent analyses are sensitive to missing data. Based on 559 samples and 47 target genes, the reference genes ASH and HDAC1 were selected using the same criteria as for pipefish females. From the geometrical mean of the two reference genes −ΔCt values were calculated to quantify relative gene expression.

*Origin salinity* (*high* or *low*), *acclimation salinity* (high or low) and *developmental salinity* (high or low) were defined as fixed factors, whereas *sampling site* (Flens, Fehm, Falck, Salz, RuegN or RuegS) was nested within *origin salinity*. A Permutational Multivariate Analysis of Variance (PERMANOVA) was applied to gene expression (–ΔCt values) of all samples and target genes for each factor and every interaction of the fixed factors. The PERMANOVA (package ‘vegan’, function ‘adonis’ in R (Oksanen, Blanchet et al. 2019)) was based on Euclidean distance matrixes with 1000 permutations (Beemelmanns and Roth 2016). A post-hoc analysis of variance (ANOVA) for every gene was applied; though, to account for multiple testing, only factors and factor interactions identified as significant by the PERMANOVA were considered. To visualize similarity/dissimilarity in gene expression among treatment groups, we performed PCAs (package: ‘ade4’, function: ‘dudi.pca’ and ‘s.class’(Dray and Dufour 2007)). To visualize significant differential gene expression among groups in heatmaps (package: ‘NMF’, function ‘aheatmap’), −ΔΔCt values for each gene were calculated as follows (Yuan, Reed et al. 2006):

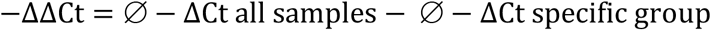

#### 2.6.3 Mortality of juveniles within the first 10 days and post-infection

Ten days post-hatch endpoint mortality of juveniles was analyzed as a ratio of “alive” vs “dead” fish using a generalized linear model (package: lme4, function: glm) with binomial error and the following fixed factors: *Origin salinity* (*High* or *Low*), *acclimation salinity* (High or Low) and *developmental salinity* (high or low) and the *sampling site* nested in *origin salinity*. Significance was tested using ANOVA type two partial sums of squares, and models were simplified using Akaike information criterion (AIC) (Akaike, 1976). Post-hoc tests were carried out using Tukey’s honest significant difference (TukeyHSD, package: multcomp, function: glht (Hothorn, Bretz et al. 2020)).Endpoint mortality of juveniles used in the infection experiment was analyzed as described above including *infection treatment* (control (c), sea water injection (sw), *Vibrio* 7 PSU (v7) and *Vibrio* 15 PSU (v15) injection) as an additional factor.

## 3. Results

### 3.1 Pipefish population structure

Allele frequencies obtained at 11 microsatellite loci of pipefish sampled at six sampling sites along the German Baltic Sea coastline indicated gene flow or recently isolated populations with no or very little divergence on neutral genetic markers. The findings are based on a Bayesian clustering method using the software STRUCTURE (Figure 2), global fixation index (F_ST_) of 0.002 and pairwise F_ST_ (S 5) Overall, the pairwise F_ST_ were low for all comparisons and with the exception of Flackenstein-Fehmarn (F_ST_ = 0.024) and Falckenstein-Flensburg (F_ST_ = 0.016) pairwise comparisons had a F_ST_ ≤ 0.01 (S 5). Populations with no differentiation on neutral markers are an ideal setting to study local adaptation because it enables us to observe phenotypic differences caused by genes under selection rather than differences caused by drift and isolation.

**Figure 2:**
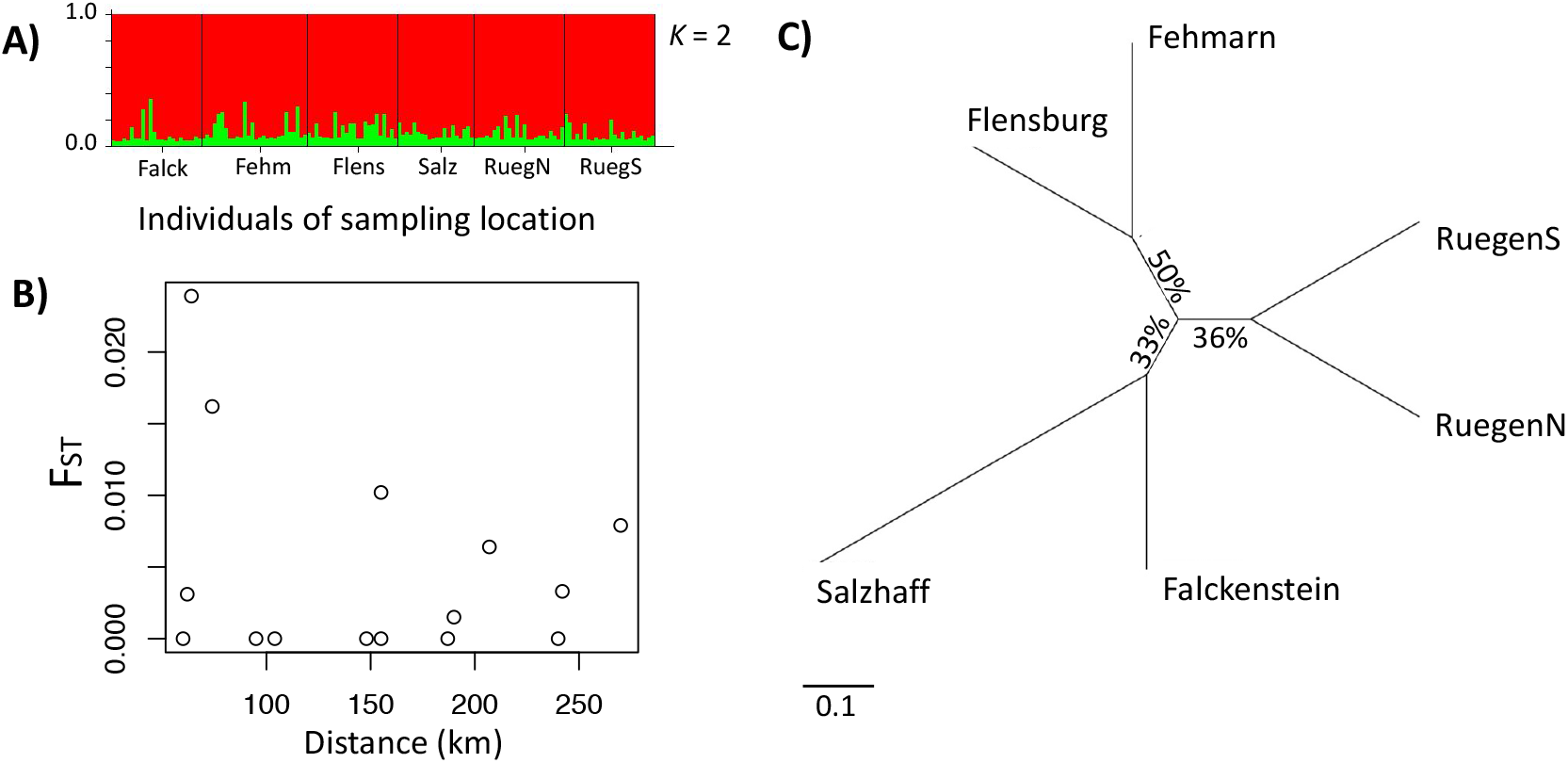
Population structure of pipefish in the southern Baltic Sea. **A:** STRUCTURE software results based on 11 microsatellite loci. Each individual is represented by a vertical line, which is colored according to the assigned groups (*K* = 2). **B:** Plotting pairwise FSTs (y-axis) against distance between sampling sampling sites (x-axis) does not reveal isolation by distance (waterway). **C:** Unrooted phylogeographic PHYLIP tree. Distances indicate the relative divergence of microsatellite loci in pipefish between sampling sites. Scale bar represents nucleotide substitutions per site.

### 3.2 Life history traits & fungus infection of parental generation

#### 3.2.1 Pipefish adults from low saline environment have a smaller body size

We found an interaction in the total length of adult pipefish between *origin salinity* and *acclimation salinity* (ANOVA F_1,320_ = 7.4, p < 0.01) indicating that parental *acclimation salinity* negatively affects growth of adult pipefish depending on the *origin salinity.* There was a trend that adults from high *origin salinity* grew slower at low *acclimation salinity* compared to high *acclimation salinity* (Tukey HSD, *H*H - *H*L: p = 0.085; S 6b), whereas *acclimation salinity* did not affect size of pipefish from low *origin salinity* (Tukey HSD, *L*L - *L*H: p = 0.535). Furthermore, all pairwise comparisons suggest that pipefish from high *origin salinity* are in general larger than pipefish from low *origin salinity* (Tukey HSD, *L*L *H*H: p < 0.001, *L*L - *H*H: p < 0.001, *L*H - *H*L: p < 0.001). The significant factor *sampling site*, which was nested in *origin salinity* (ANOVA F_4,320_ = 11.2, p < 0.01) indicates that individuals from Salzhaff were larger compared to individuals from Ruegen North and Ruegen South but did not differ from pipefish caught at the high *origin salinity* (Tukey HSD, Salz - RuegN: p < 0.001; Salz - RuegS: p < 0.001; S 6c).

The correlation between the salinity at the sampling site and the size of the adults, i.e. length (Figure 3 B) and weight (Figure S6) suggest that pipefish from low *origin salinity* were smaller. The total length and weight in these plots are not corrected for age, which has not been assessed. However, pipefish are usually all in the same age cohort when they are caught in spring. Most of them were born the summer before and reached sexual maturity around the time of catching.

**Figure 3:**
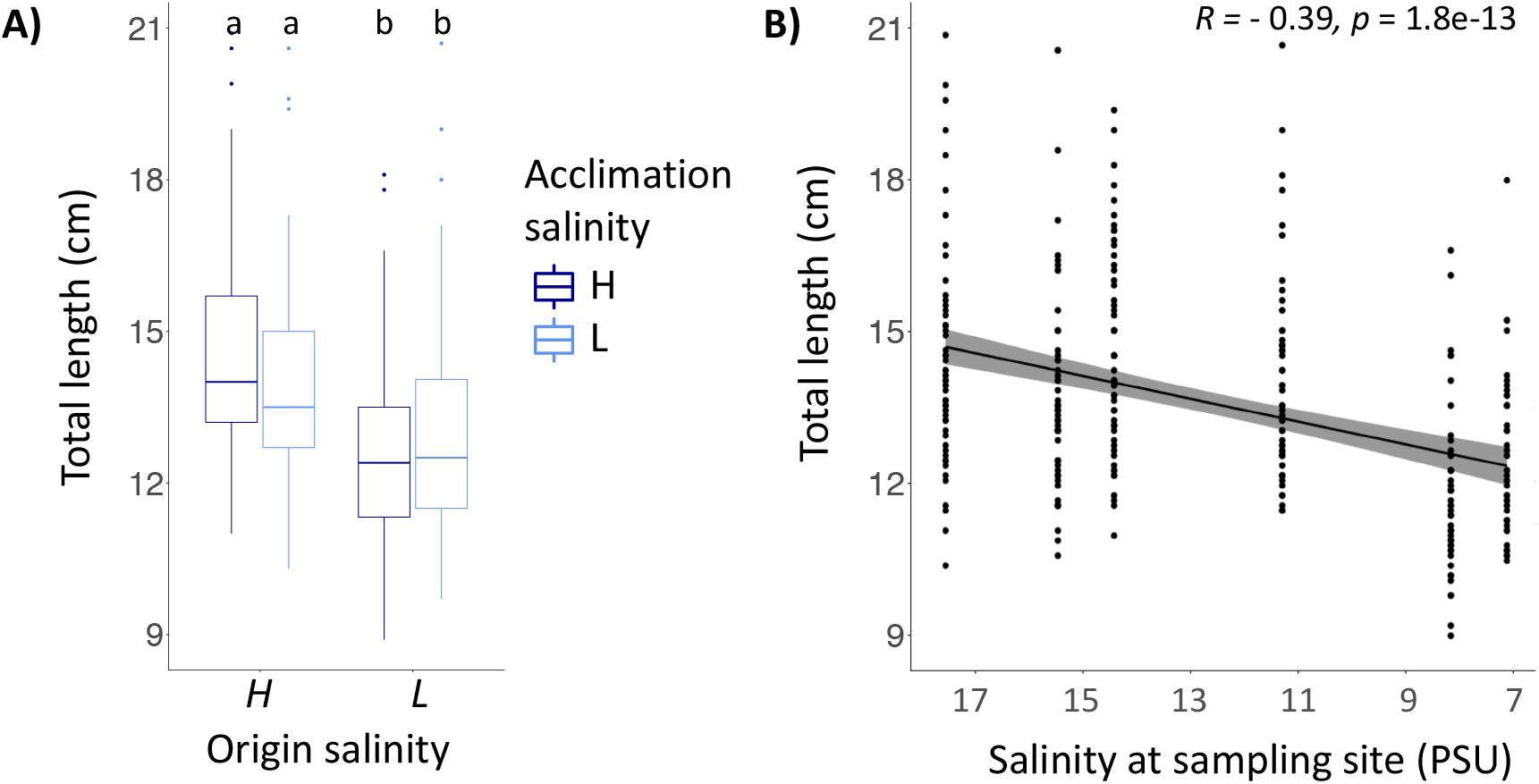
Body length of adult pipefish (A) correlated with salinity at the sampling site (B) **A:** The total length of pipefish is shown for both salinity categories (high - 15 PSU, low −7 PSU). Colors indicate whether the parental generation was acclimated to a high (H, dark blue) or to low saline environment (L, light blue). **B:** The total length of pipefish after acclimation correlated with the salinity measured at the sampling site using spearman rank correlation. Grey bar indicates 95% confidence interval.

#### 3.2.2 Pipefish from high saline environments were more susceptible to fungus infection when exposed to low saline conditions

Visible fungus infections of the brood pouch occurred in almost half of the pipefish males (47%) caught at high *origin salinity* and kept at low *acclimation salinity* (Figure 4). Fungus infections ranged from mild infections in the brood pouch not affecting clutch size to a complete loss of the offspring. Males from a high *origin salinity* that remained at high *acclimation salinity* as well as males from the low *origin salinity* had no symptoms of fungus infection regardless of the *acclimation salinity*.

**Figure 4:**
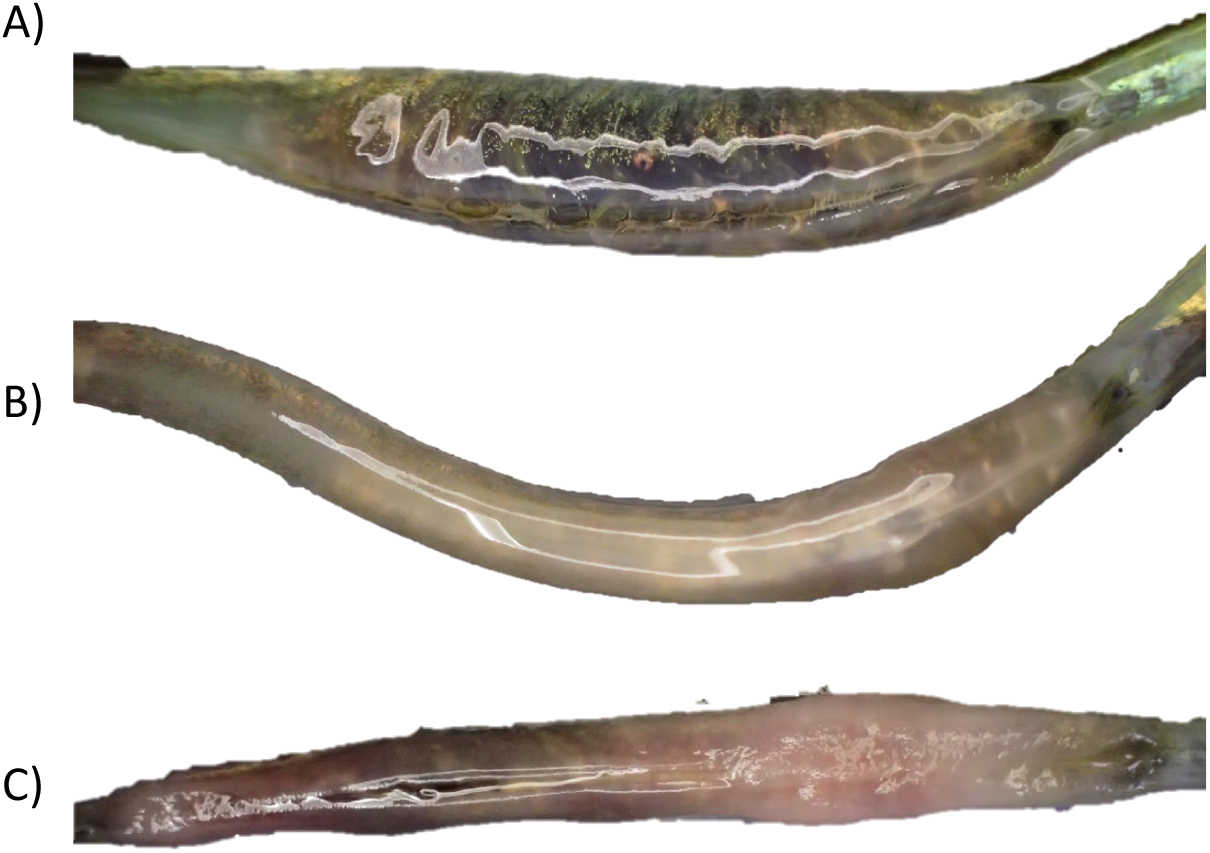
Brood pouch fungus infections were present in 47 % of fathers from high salinity origin kept at low acclimation salinity. Pipefish from low *origin salinity* independent of *acclimation salinity* and pipefish from high *origin salinity* kept at high *acclimation salinity* did not show any signs of fungus infection (A). In males caught at high *origin salinity* and kept at low *acclimation salinity* fungi infections ranged from mild (B) to extreme (C) resulting in the complete loss of eggs and offspring.

#### 3.2.3 Clutch size

Males from a high *origin salinity* kept at high *acclimation salinity* had the largest clutch size (*H*H, mean ± s.d., 41.8, ± 23.4, Tukey TSD; S 7b) followed by males from low *origin salinity* kept at high salinity (*L*H, 27.8 ± 13.2) or low salinity (*L*L, 25.2 ± 16.9), which corresponds to the lower body size at low salinity (Figure 3). Pipefish from high *origin salinity* exposed to low *acclimation salinity* were frequently infected by a brood pouch fungus which reduced the clutch size (*H*L 19.4 ± 15.2; Tukey TSD, *H*L-*H*H, t = - 4.8, p < 0.001;S 7b). In contrast, *acclimation salinity* did not affect clutch size of parents from low *origin salinity* (Tukey TSD, *L*L-*L*H, t = - 0.4, p = 0.976; S 7b) This divergent patterns caused a o*rigin salinity*:*acclimation* salinity interaction (ANOVA F_1,109_ = 9.0, p = 0.003; S 7a). Higher clutch size, was in general driven by a larger total body length of male pipefish (Spearman rank correlation, R = 0.43, p < 0.001; Figure 5)

**Figure 5:**
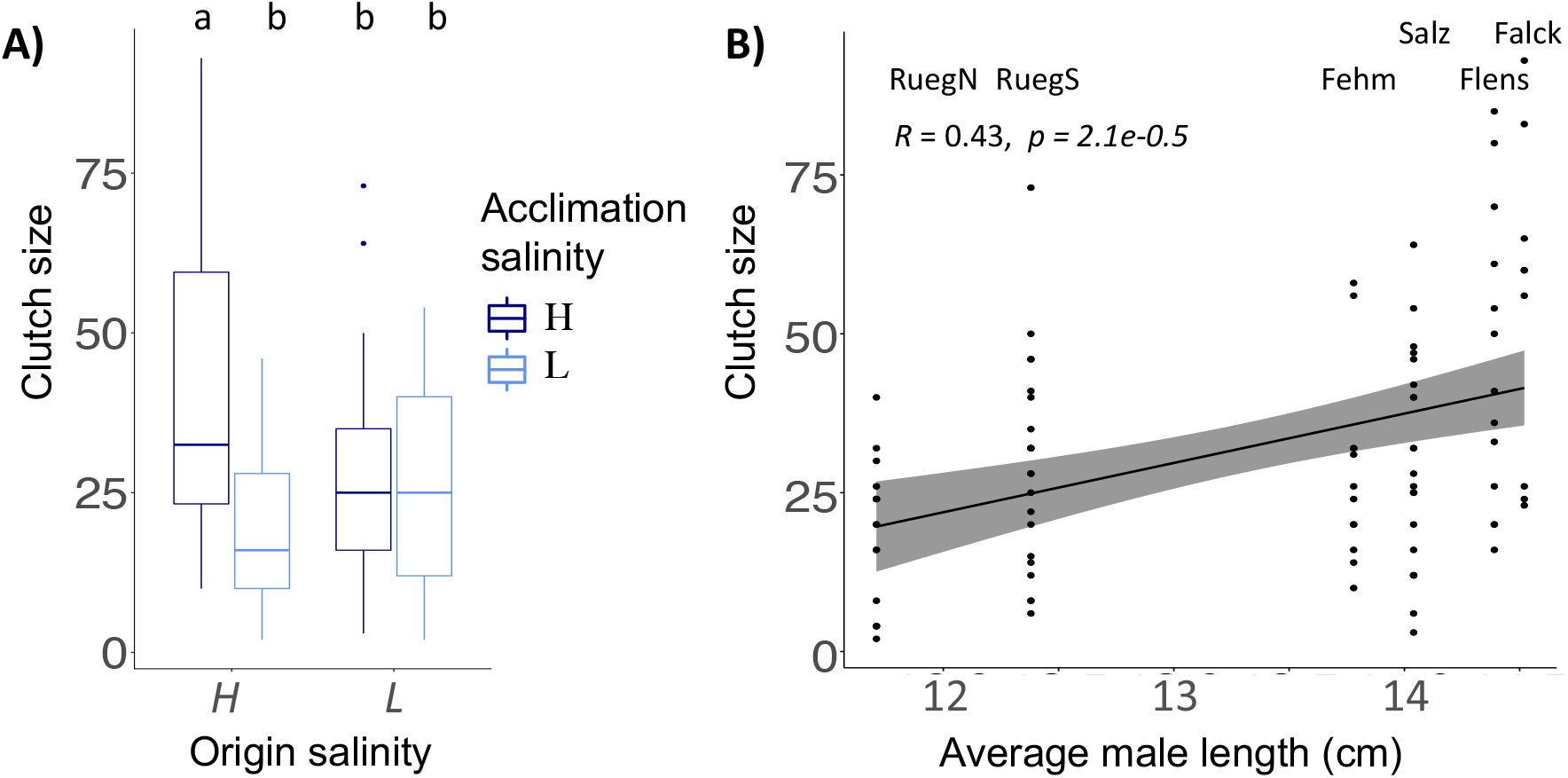
Parental life history traits. **A:** Number of juveniles (clutch size) is shown for pipefish with high (*H*) or low (*L*) *salinity origin*. Color indicates *acclimation salinity* in the lab (H: 15 PSU - dark blue and L: 7 PSU - light blue). Different letters indicate significant differences (Tukey’s HSD, p < 0.05). **B:** Clutch size correlated positively with the average size of male pipefish from each sampling site. The *sampling sites* are presented above the data points. The grey bar along the regression line indicates the 95% confidence interval. A more detailed figure visualizing the differences between the single origins can be found in the supplement (S 7c).

#### 3.2.4 Two immune genes are upregulated in females from a high origin salinity

*Origin salinity* had an impact on the expression of immune genes in female pipefish (PERMANOVA, *immune* F_1,146_ = 3.1, p = 0.010; S 8), in particular of the innate immune system (PERMANOVA, *innate* F_1,146_ = 4.0, p = 0.002). *Lectin protein type I* (*Lecpt1*), a pathogen recognition receptor, and *chemokine 7* (*ck7*), a gene encoding a protein responsible for chemotaxis in blood cells, were upregulated in pipefish from high *origin salinity* in contrast to low *origin salinity* females. Low *acclimation salinity* caused a slight upregulation in the expression of histone modification gene *histone deacetylase 6-like* (*hdac6*) (PERMANOVA, *silencing* F_1,146_ = 2.9, p = 0.044).

#### 3.2.5 Juveniles from low origin salinity parents have higher survival rates and are smaller

In the first ten days after hatching, patterns of juvenile survival suggest an *origin salinity*:*acclimation salinity* interaction (GLM, *χ*^2^_1_ = 6.1, p = 0.031; S 9a). Whereas, survival of juveniles from parents continuously exposed to the same salinity (Tukey HSD, *L*L - *H*H: z = −3.4, p = 0.783; S 9b) or non-matching *origin* and *acclimation salinity* did not differ (*L*L *L*H: z = −4.7, p = 0.842; *H*H - *H*L: z = −2.1, p = 0.148), juveniles from high *origin salinity* parents exposed to high *acclimation salinity* in the lab (*L*L) had higher survival rates compared to juveniles from high *origin salinity* exposed to low *acclimation salinity* (Tukey HSD, *L*L - *H*L: z = −3.4, p = 0.043; S 9b). The *origin salinity:sampling site* effect suggest that patterns at single sampling sites differ. In particular, Flensburg offspring exposed to high *developmental salinity* had reduced survival rates, when parents were acclimated to low instead of high salinity (Tukey’s HSD; Hh - Lh: z = −4.4, p = 0.046; S 9b). Following the same pattern of non-matching *acclimation salinity*, Ruegen South offspring exposed to low *developmental salinity* had reduced survival, when parents were kept at high *acclimation salinity* (Tukey HSD; Hl - Ll: z = 5.8, p < 0.001). This suggests that exposure of parents to novel salinities can negatively impact juvenile survival when juveniles experience salinity conditions, which did not match parental *acclimation salinity*.

Overall higher survival at high *developmental salinity* compared to low *developmental salinity* (*developmental salinity*, GLM, *χ*^2^_1_ = 192.8, p = 0.031; Figure 6; S 9a) indicates that low salinity imposes a stress on pipefish juveniles regardless of the *salinity origin*. An exception are juveniles from Ruegen North (*sampling site*, GLM, *χ*^2^_4_ = 24.1, p < 0.001; S 9a) where juvenile survival was not affected by *developmental salinity* (GLM, *χ*^2^_1_ = 0.1, p = 0.766)

**Figure 6:**
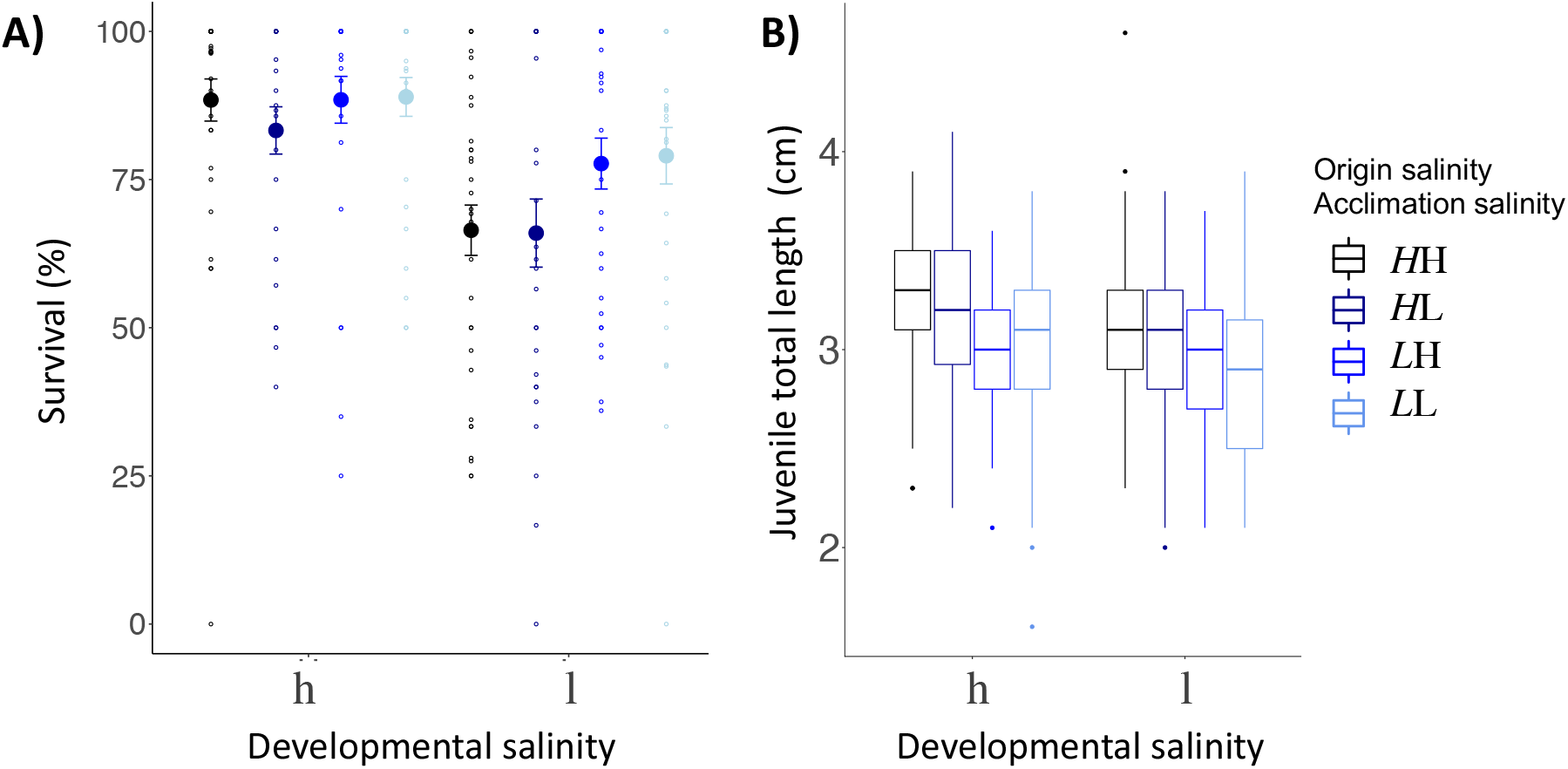
Juvenile life history parameters (A) survival rate (%), (B) body length (cm) **A:** The percentage of juveniles surviving the first 10 days post-hatch (%) are plotted on the y-axis. The x-axis indicates the *developmental salinity* after hatching (h – high/15 PSU, l – low/7 PSU). Italic letters and colors specify the *origin salinity* of the parental generation (*H*: 15 PSU, black & dark blue; *L*: 7 PSU blue & light blue). The 2nd letter indicates the *acclimation salinity* in the lab (H: 15 PSU, black and blue; L: 7 PSU, dark & light blue). **B:** Juvenile size (cm) 10 days post hatch is plotted on the y-axis. Labelling and color code correspond to Panel A.

Ten days after hatching, juveniles from high *origin salinity* were larger (3.18 cm ± 0.37, n = 408) than juveniles from low *origin salinity* sampling sites (2.95 cm ± 0.37, n = 405) (*origin salinity*, ANOVA F_1,782_ = 86.2, p < 0.001; S 10). While *acclimation salinity*, i.e. mating and male pregnancy, had no effect on size of juveniles (*acclimation salinity*, ANOVA F_1,782_ = 2.2, p < 0.136), low *developmental salinity* reduced offspring size (*developmental salinity*, ANOVA F_1,782_ = 17.4, p < 0.001) suggesting are stressful for pipefish offspring and reduce growth rates.

#### 3.2.6 Juvenile survival is reduced after injections and at low salinity

Ten days post hatch, juvenile pipefish were challenged either with *Vibrio alginolyticus* bacteria evolved at 15 PSU, 7 PSU, autoclaved seawater (sham injection) or not treated at all (control) and survival was measured six days post infection, i.e. approximately 16 days post hatch. Non-challenged control groups had the highest survival rates (Mean ± s.d.; 83.0% ± 32.2; Figure 7). The injection itself decreased survival of juveniles by at least 10% in all salinity treatments combined, regardless whether seawater (66.9% ± 38.2), *Vibrio* evolved at 15 PSU (73.0% ± 36.8) or 7 PSU (66.7% ± 38.6) was administered. *Vibrio* strains evolved at 15 PSU caused a higher mortality in juveniles from high *origin salinity* regardless of *acclimation salinity* compared to juveniles from low *origin salinity* with low parental *acclimation salinity* (*origin salinity* x *acclimation salinity* x *treatment,* GLM, *χ*^2^_1_ = 13.0, p = 0.005; S 11; Tukey HSD; *L*Lv15 - *H*Hv15: z = −3.4, p = 0.046; *L*Lv15 - *H*Lv15: z = −3.5, p = 0.038). When fathers from low *origin salinity* were exposed to high *acclimation salinity* these positive effects on offspring survival were lost (Tukey’s HSD, *L*Hv15 - *H*Hv15 z = −1.5, p = 0.971). This suggests that mis-matching salinity levels between the parental and juvenile generation can lead to reduced survival rates and that juveniles of fathers from low salinity levels have higher survival rates compared to juveniles of fathers from high salinity.

**Figure 7:**
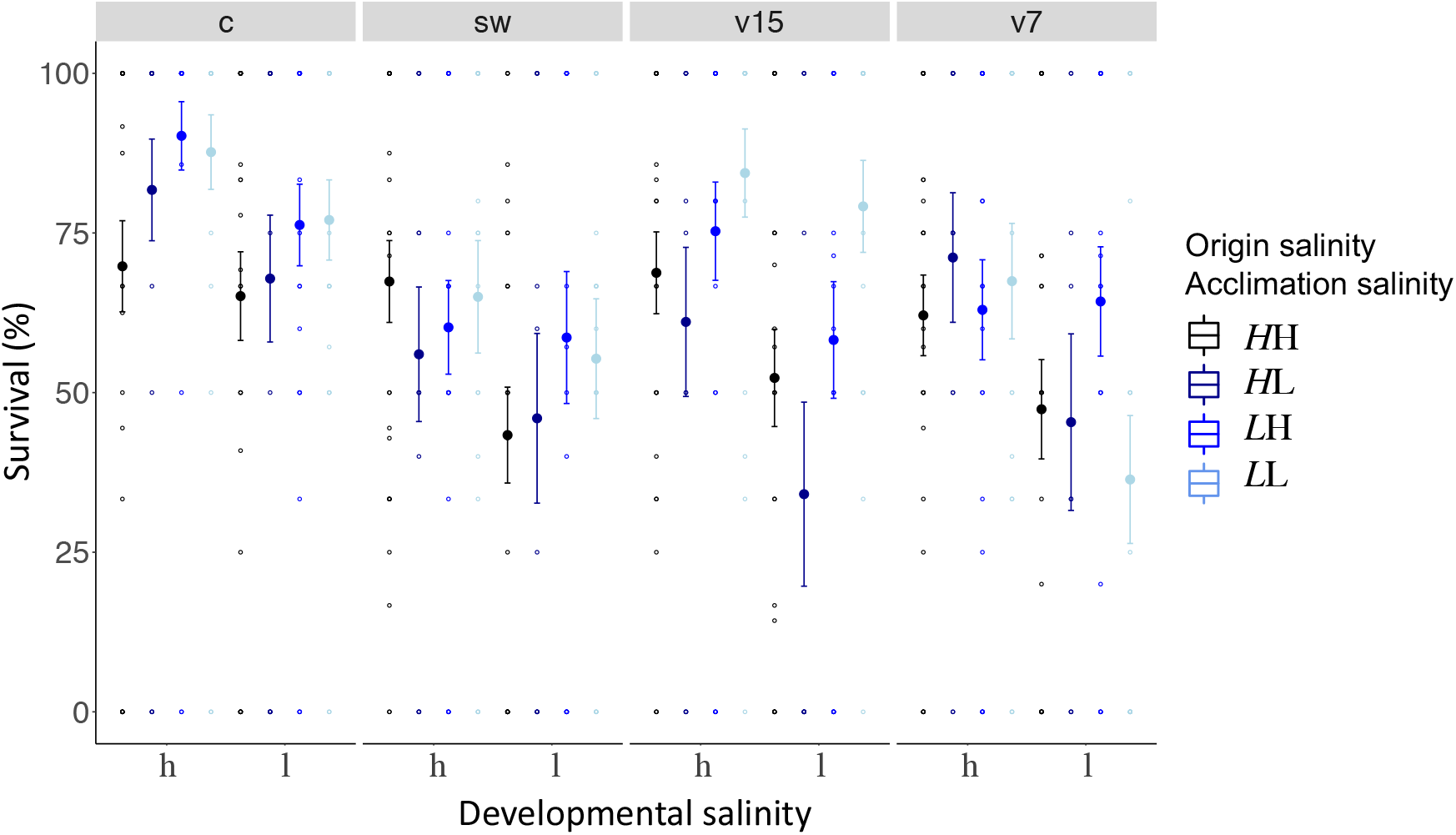
Juvenile survival six days post infection. Juvenile survival six days post infection is plotted on the y-axis. The x-axis indicates the *developmental salinity* (h - high/15 PSU, l - low/7PSU). Italic letters and colors specify the *origin salinity* of the parental generation (*H*: 15 PSU, black & dark blue; *L*: 7 PSU blue & light blue). The 2nd letter and colors indicate the *acclimation salinity* in the lab (H: 15 PSU, black and blue; L: 7 PSU, dark & light blue). Each *treatment* is represented by one panel, i.e. control (c) or injection with seawater (sw), *Vibrio* strain evolved at 15 PSU (v15), or at 7 PSU (v7).

Juvenile survival was in general higher in high *developmental salinity* conditions (71.1% ± 36.7) compared to low *developmental salinity* conditions (58.6% ± 36.7) (GLM, *developmental salinity*, *χ*^2^_1_ = 40.2, p = 0.031) suggesting that low salinity levels are a stressful environment for pipefish development. An adaptation to low salinity may result in an increased fitness as juveniles from low *origin salinity* fathers (69.8% ± 40.1) had in general a higher survival rate compared to juveniles from high *origin salinity* fathers (60.6% ±37.7).

An effect of *sampling site* nested in *origin salinity* (GLM, *χ*^2^_4_ = 39.5, p < 0.001; S 11) indicates that survival patterns for each sampling site within the *origin salinity* categories are diverse. The statistical diversity may be a result of the high variation in survival rates within a single treatment, which sometimes ranged from 0 - 100%. Combining the survival rates of all three sample sites of one *origin salinity* resulted in more robust and conclusive results.

#### 3.2.7 Matching parental acclimation and juvenile developmental salinity results in similar juvenile gene expression patterns of adaptive immune genes

An *origin salinity* effect indicates that gene expression of juvenile differs depending on the salinity they originate from (PERMANOVA, *all genes; origin salinity* F_1,523_ = 4.2, p = 0.003). Such signs of genetic adaptation were found in genes associated with the innate immune system (PERMANOVA, *innate,* F_1,523_ = 9.2, p = 0.001) and with osmoregulation (PERMANOVA, *osmo*, F_1,523_ = 4.1, p = 0.003; Figure 8, Table 2). Single ANOVAs suggested that this effect was driven by five genes. Whereas the pathogen reception recognition gene *lectin protein type II* (*lectpt2*) was higher expressed in parents from low *origin salinity*, the expression of the following genes was upregulated in juveniles from high *origin salinity* parents: *immunoglobulin light chain* (*IgM*; pathogen recognition), *heat shock protein 70 kDa* (*hsp70;* osmotic stress response); *voltage gated potassium channel* (*kcnh8;* cell volume regulation)*, prolactin (prl;* ion uptake promotion and ion secretion inhibition). The genetically induced upregulation of osmoregulatory genes suggests an adaptation to low salinity levels.

**Figure 8:**
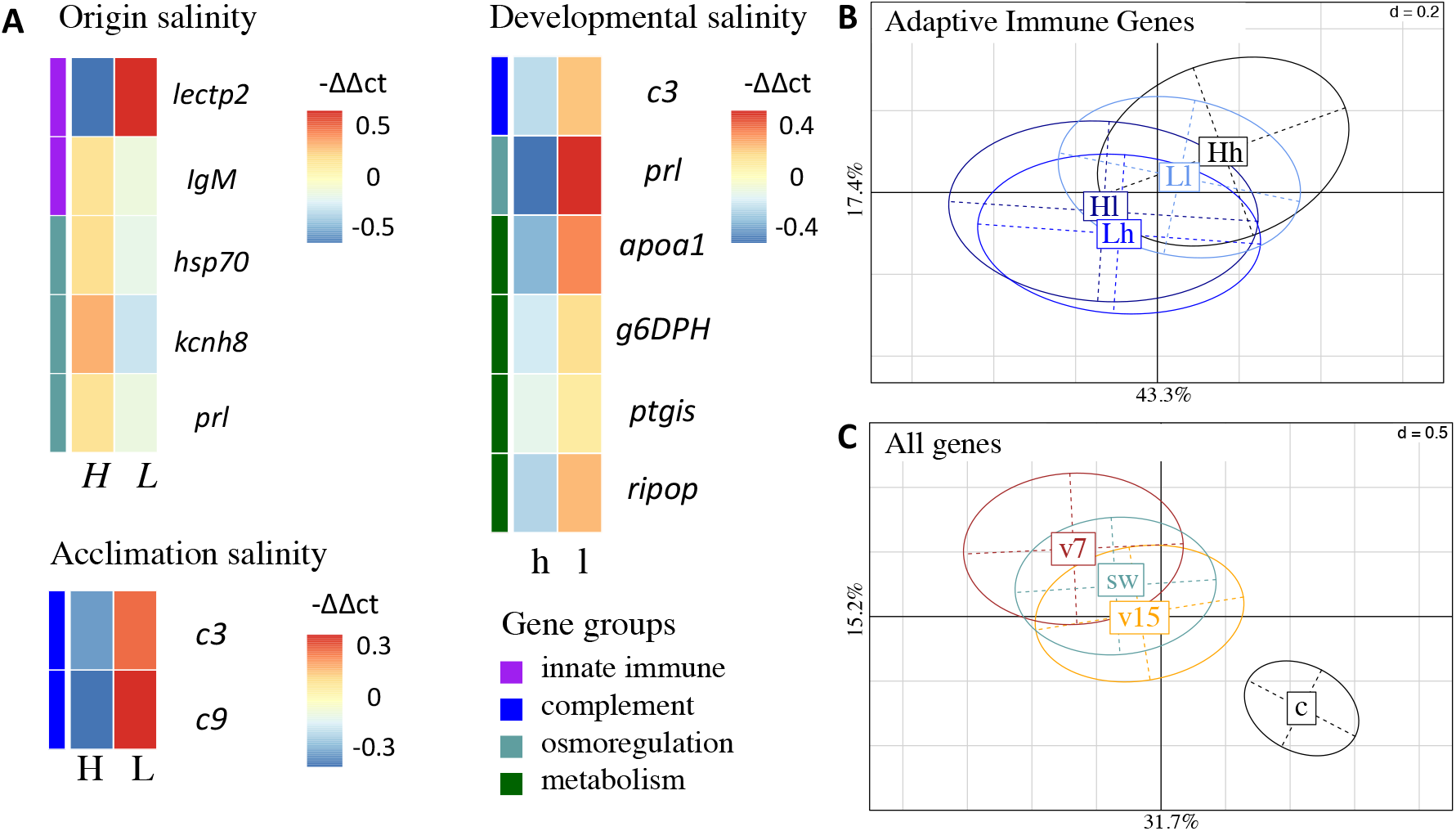
Gene expression patterns of juveniles. **A:** Non-hierarchical gene expression heatmap for genes showing differential expression (−ΔΔCt) in response to *origin salinity*, *acclimation salinity* or *developmental salinity*. Genes are sorted by gene groups which are assigned with different colors (purple: innate immune system, blue: complement system, turquoise: osmoregulation, green: metabolism). **B:** Principal component analysis (PC1: 43.3%; PC2: 17.4%) of adaptive immune genes for significant interaction between parental *acclimation salinity* (H – high (15 PSU), L – low (7 PSU)) and *developmental salinity* (h, l). Four out of seven adaptive immune gene were upregulated (S 4), when parental *acclimation salinity* and juvenile *developmental salinity* did not match, i.e. Hl and Lh, compared to a matching *acclimation* and *developmental salinity*, i.e. Hh and Ll (PERMANOVA, *adaptive* F1,523 = 2.6, p = 0.034). The confidence ellipse explains 20 % of the variability. **C:** Principal component analysis (PC1: 31.7%; PC2: 15.2%) of gene expression patterns caused by juvenile *treatment*, i.e. injection with *Vibrio alginolyticus* evolved at 15 PSU (v15, orange), *V. alginolyticus* evolved at 7 PSU (v7, brown), sham injection with seawater (sw, bluegreen) or untreated control (c, black). The confidence ellipse explains 20 % of the variability

**Table 2:** PERMANOVA results of juvenile gene expression: A PERMANOVA was applied to gene expression (−ΔCt values) of all 558 samples for all genes (47), immune genes comprised of the innate, adaptive and complement genes, as well as genes involved in metabolism, osmoregulation and epigenetics, e.g. methylation or histone modification. Results are based on Euclidian distance matrices with 1000 permutations. Significant *p*-values are in bold.

|  |  | All genes |  | Immune |  | Innate |  | Adaptive |  | Complement |  | Metabolism |  | Osmoreg |  | Epigenetics |  |
| --- | --- | --- | --- | --- | --- | --- | --- | --- | --- | --- | --- | --- | --- | --- | --- | --- | --- |
|  | df | F<br>Model | Pf(>F) | F<br>Model | Pf(>F) | F<br>Model | Pf(>F) | F<br>Model | Pf(>F) | F<br>Model | Pf(>F) | F<br>Model | Pf(>F) | F<br>Model | Pf(>F) | F<br>Model | Pf(>F) |
| Origin salinity (Origin sal) | 1 | **4.2** | **0.003\*** | 5.2 | **0.001\*** | **9.2** | **0.001\*** | 1.4 | 0.21 | 2.3 | 0.098 | 1.5 | 0.203 | **4.1** | **0.003\*** | 0.7 | 0.508 |
| Acclimation salinity Acclim sal) | 1 | 1.5 | 0.15 | 1.6 | 0.129 | 1.2 | 0.277 | 0.5 | 0.715 | **4.6** | **0.014\*** | 1.1 | 0.326 | 1 | 0.411 | 2.6 | 0.578 |
| Developmental salinity (Devo sal) | 1 | **3.2** | **0.005\*** | **2.5** | **0.034\*** | 1.7 | 0.136 | 1.3 | 0.234 | **3.3** | **0.028\*** | **4.2** | **0.020\*** | **6.4** | **0.001\*** | 1.2 | 0.298 |
| Treatment (Treat) | 3 | **11.3** | **0.001\*** | **13.9** | **0.001** | **17.5** | **0.001\*** | **16.8** | **0.001\*** | **6.3** | **0.001\*** | **2.5** | **0.025\*** | **9.1** | **0.001\*** | **3.2** | **0.005\*** |
| Origin sal:Acclim sal | 1 | 0.5 | 0.858 | 0.4 | 0.93 | 0.3 | 0.91 | 0.7 | 0.582 | 0.6 | 0.535 | 0.2 | 0.908 | 0.8 | 0.525 | 1.6 | 0.182 |
| Origin sal:Devo sal | 1 | 0.6 | 0.73 | 0.6 | 0.783 | 0.6 | 0.663 | 0.6 | 0.642 | 0.3 | 0.74 | 0.3 | 0.798 | 1.2 | 0.265 | 0.5 | 0.672 |
| Acclim sal:Devo sal | 1 | 1.6 | 0.141 | 1.8 | 0.088 | 2.1 | 0.639 | **2.6** | **0.034\*** | 1.3 | 0.26 | 0.8 | 0.436 | 1.2 | 0.311 | 2.2 | 0.093 |
| Origin sal:Treat | 3 | 1.1 | 0.289 | 1.2 | 0.255 | 0.9 | 0.502 | 0.6 | 0.782 | 1.5 | 0.19 | 2.1 | **0.037\*** | 0.7 | 0.796 | 0.6 | 0.762 |
| Acclim sal:Treat | 3 | 0.6 | 0.933 | 0.4 | 0.995 | 0.3 | 0.999 | 0.8 | 0.633 | 0.3 | 0.947 | 0.9 | 0.503 | 0.9 | 0.526 | 0.5 | 0.881 |
| Devo sal:Treat | 3 | 0.8 | 0.675 | 0.9 | 0.603 | 0.8 | 0.671 | 1.1 | 0.336 | 0.6 | 0.722 | 0.9 | 0.453 | 0.8 | 0.678 | 0.6 | 0.815 |
| Origin sal:sampling site | 4 | **2.9** | **0.001\*** | **3.1** | **0.001\*** | **3.5** | **0.001\*** | **3.2** | **0.001\*** | **2.2** | **0.036\*** | **2.3** | **0.023\*** | **2.1** | **0.005\*** | **3.7** | **0.003\*** |
| Origin sal:Acclim sal:Devo sal | 1 | 1.2 | 0.251 | 1 | 0.372 | 1.3 | 0.221 | 1.1 | 0.354 | 0.3 | 0.809 | 1.4 | 0.214 | 0.7 | 0.633 | 2.2 | 0.089 |
| Origin sal:Acclim sal:Treat | 3 | 0.8 | 0.725 | 0.8 | 0.705 | 0.7 | 0.805 | 0.9 | 0.513 | 1 | 0.42 | 0.9 | 0.498 | 0.8 | 0.669 | 0.9 | 0.482 |
| Origin sal:Devo sal:Treat | 3 | 0.9 | 0.604 | 0.6 | 0.877 | 0.5 | 0.917 | 0.4 | 0.953 | 0.6 | 0.72 | 1.2 | 0.28 | 1.1 | 0.307 | 0.9 | 0.465 |
| Acclim sal:Devo sal:Treat | 3 | 0.9 | 0.534 | 0.9 | 0.582 | 0.8 | 0.618 | 1.3 | 0.202 | 1.1 | 0.37 | 0.7 | 0.628 | 0.8 | 0.665 | 0.9 | 0.541 |
| Origin sal:Aacclim sal:Devo sal:<br>Treat | 3 | 0.6 | 0.933 | 0.5 | 0.983 | 0.5 | 0.956 | 0.5 | 0.912 | 0.4 | 0.918 | 0.6 | 0.718 | 0.9 | 0.507 | 0.7 | 0.664 |
| Residuals | 523 |  |  |  |  |  |  |  |  |  |  |  |  |  |  |  |  |
| Total | 558 |  |  |  |  |  |  |  |  |  |  |  |  |  |  |  |  |

Juvenile gene expression did not provide further evidence for genetic adaptation, tested as an interaction between *origin salinity* and *acclimation salinity* (PERMANOVA, *all genes* F_1,523_ = 0.5, P = 0.858). Trans-generational plasticity could also not be detected in the interaction between *acclimation salinity* and *developmental salinity* (PERMANOVA, *all genes* F_1,523_ = 0.6, P = 0.730).

Exposing parents to low *acclimation salinity* led to an expression induction of two genes in juveniles (PERMANOVA, *complement; acclimation salinity* F_1,523_ = 4.6, p = 0.014). Both genes are associated with the complement system*: Complement component 3* (c3, complement system activation) and *Complement component 9* (*c9*, membrane attack complex). In addition to this parental effect a trans-generational effect was observed as a *parental acclimation salinity* : *developmental salinity* interaction effect on adaptive immune gene expression (PERMANOVA, *adaptive* F_1,523_ = 2.6, p = 0.034). Gene expression was lower in four out of seven adaptive immune genes when *acclimation salinity* and *developmental salinity* were matching compared to non-matching conditions (S 12): *Human immunodeficiency virus type l enhancer* 2 (*hivep2*, transcription factor, MHC enhancer binding) and *3* (*hivep3;* transcription factor, MHC enhancer binding*), B-cell receptor-associated protein (becell.rap31*, T- and B-cell regulation activity) and *immunoglobulin light chain* (*igM;* antigen/pathogen recognition). The reduction in gene expression of immune genes can hint at a reduced stress level in offspring fish when parents are acclimated to the same salinity as their offspring.

Developmental plasticity allows juveniles to quickly respond to present salinity levels. Low *developmental salinity* resulted in higher expression of six gens. *Complement component 3 (c3)* is involved in the complement system (PERMANOVA, *comple; developmental salinity* F_1,523_ = 4.6, p = 0.014; Figure 8, Table 2), *prolactin* (*prl*, ion uptake promotion and ion secretion inhibition) is associated with osmoregulation (PERMANOVA, *osmo; developmental salinity* F_1,523_ = 3.3, p = 0.028) and *apolipoprotein A1* (*apoa1,* antimicrobial activity*), glucose 6 phosphate dehydrogenase* (*g6DPH*, pentose phosphate pathway)*, prostaglandin I2 Synthase* (*ptgis,* Lipid and fatty acid metabolism), *ribosomal protein* (*ripop*, Translation process) are related to the metabolism (PERMANOVA, *meta; developmental salinity* F_1,523_ = 4.6, p = 0.014).

Finally, we wanted to test whether genetic background, i.e. *origin salinity*, *acclimation salinity* of the parents and the *developmental salinity* influenced the ability of juveniles to cope with infections of the opportunistic pathogen *V. alginolyticus*, which evolved in the lab at either 15 or 7 PSU. However, we found no interaction between any salinity regime of parents or juveniles interacting with gene expression after juvenile infection. An injection, regardless of the component, i.e. autoclaved seawater, *V. alginolyticus* evolved at 15 PSU or 7 PSU caused similar changes in gene expression patterns that could only be differentiated from the untreated control group. In 24 genes injections caused a higher gene expression (PERMANOVA, *all genes;* F_3,523_ = 11.3, p = 0.001; S 13), including genes from all groups. In five genes, injections caused a lower gene expression compare to the control group. Using post-hoc tests on ANOVAs of single genes, we found no differences in gene expression between the three injection treatments.

## 4. Discussion

In the present study we investigated the role of genetic adaptation and phenotypic plasticity as well as their interaction on the ability of the broad-nosed pipefish *Syngnathus typhle* to cope with changes in salinity levels. *S. typhle* is a marine teleost, which originally invaded from the North Sea into the Baltic Sea (Wilson and Veraguth 2010). The brackish salinity environment in the Baltic Sea imposes osmoregulatory stress on marine animals and is thus assumed to be an important driver for genetic divergence and adaptation to local condition (Berg, Jentoft et al. 2015, Guo, DeFaveri et al. 2015, Johannesson, Le Moan et al. 2020). Here, we focused on six pipefish populations from the Baltic Sea, out of which three originated from a rather high saline environment (14 - 17 PSU), and three from a low saline environment (7 - 11 PSU). By taking the two salinity regimes into account, our experiment permitted to test both for local adaptation and for phenotypic plastic acclimation to different salinities.

*S. typhle* caught in the Baltic Sea high salinity environments (14 - 17 PSU) were smaller (14.2 cm) than those populating the marine realm with more than 28 PSU (mean size animals caught between 28 and 36 PSU: 18.7 cm (Rispoli and Wilson 2008) or 15.5 cm (Gurkan and Taskavak 2007), but larger than those sampled in Baltic Sea low salinity environments (mean size in this study: 12.8 cm; pipefish sampled at 5.5 PSU around Askö (Sweden): 14.5 cm (Rispoli and Wilson 2008). This suggests that osmoregulation is costly (Rolfe and Brown 1997, Boeuf and Payan 2001) and that the negative impact of low salinity can potentially not be fully compensated through local adaptation This implies that trade-offs for osmoregulation reduce growth rates, which ultimately correspond to a decreased fitness. Studies of other marine teleosts that originated from fully marine environments, e.g. sticklebacks and cod, suggested that high growth rates at intermediate salinity levels (10 - 20 PSU) are possible, especially when close or slightly above isosmotic levels (Dutil, Lambert et al. 1997, Imsland, Foss et al. 2001, Heckwolf, Meyer et al. 2018).

In this and a previous study (Nygard, Kvarnemo et al. 2019), the parental phenotype correlated with the body weight or length of the offspring. The heritability of morphological traits was suggested to be lower for ectotherms than for endotherms (Mousseau and Roff 1987). However, body size of pipefish females is known to correlate with egg size, and larger fathers were shown to give birth to embryos of an induced size (Nygard, Kvarnemo et al. 2019). To this end, both the parental body size and a resource-allocation trade off imposed by an increased energy demand for osmoregulation can explain the reduced embryonal growth in the low saline environment (Boeuf and Payan 2001). In the here presented survival experiment, juveniles from low origin salinity parents survived better compared to high origin salinity parents, independent of the parental acclimation salinity, developmental salinity and exposure to *Vibrio* bacteria. We thus suggest two alternative parental care strategies: large broad-nosed pipefish parents can invest in larger clutch and offspring size, while small parents may rather invest in survival (Nygard, Kvarnemo et al. 2019) via genetically determined gene expression patterns.

Such genetically determined gene expression patterns that are inherited from generation to generation, can be indicative signs for local adaptation (Larsen, Schulte et al. 2011, Fraser 2013, Heckwolf, Meyer et al. 2020). Females from high origin salinity had an induced baseline innate immune gene expression compared to females originating from low salinity environment. This induced innate immune gene expression pattern was inherited to their offspring: juveniles from animals caught from high saline environments generally had an induced expression of innate immune genes. We suggest that the observed induction of innate immune genes in pipefish originating from high saline origins is indicative for the existence of sufficient resources allowing to keep the innate immune response at a high baseline level. This can result in a faster and stronger and eventually more effective immune response. In contrast, pipefish from low origin salinity may rather suffer stress induced by the above stated resource allocation trade-off, which decreases the resources available for the innate immune system.

Under stress, animals are more susceptible to infections with pathogens, which may turn opportunistic pathogens into causative agents of deadly diseases (Boyett, Bourne et al. 2007, Poirier, Listmann et al. 2017, Sullivan and Neigel 2018). Furthermore, low saline environments have been suggested to select for increased pathogenic virulence, e.g. due to changes in gene expression (Hase and Barquera 2001) and biofilm formation (Dayma, Raval et al. 2015). This is in line with the observed brood pouch infections during pregnancy that massively impacted fathers adapted to a high origin salinity but exposed to low acclimation salinity. Their clutch sizes at birth were reduced and their offspring were smaller. Fathers caught at low origin salinity (*L*L & *L*H) did not show signs of brood pouch infection, which gives support for our hypothesis that these animals were locally adapt to low saline environments and the associated pathogens. An adaptation to potentially more virulent infections at low saline conditions was also found in our experiment: juveniles infected with *Vibrio* bacteria survived better when the parents were caught at low originated from low saline environments.

Juveniles are expected to have advantages when exposed to the same environment as their parents (Sunday, Calosi et al. 2014, Roth, Beemelmanns et al. 2018). An interaction of the parental acclimation salinity and the juvenile developmental salinity is generally interpreted as an indicator for trans-generational plasticity (Uller, Nakagawa et al. 2013, Heckwolf, Meyer et al. 2018). In contrast to previous experiments focusing on trans-generational plasticity and immune priming in pipefish (Beemelmanns and Roth 2016, Beemelmanns and Roth 2017, Roth and Landis 2017), the adaptive trans-generational plastic effects identified in this study were limited. Even though survival of juveniles was higher in matching parental acclimation and developmental salinity, the effect was driven by the genetic adaptation and not the parental acclimation. The same applied for juvenile growth, which was imposed both by origin salinity and by the developmental salinity, but not by acclimation salinity. However, parental acclimation shifted expression of genes involved in complement and adaptive immune systems. As such, parental acclimation to low salinity (main effect) induced the expression of genes of the complement system. Non-matching parental acclimation and developmental salinity (interaction) upregulated genes of the adaptive immune system compared to matching parental acclimation and developmental salinity. In contrast to the above discussed upregulation of innate immune genes, an upregulation of the complement and adaptive immune system is indicative for a clear response towards prevailing parasites and pathogens, due to the specificity of the adaptive immune system (Janeway 2005). The complement system links the innate to the specific adaptive immune system. Their joint induction could give evidence for a shift in the microbial pathogen community in non-matching environments to which the specific arm of the immune system has to react. However, final support would enquire the genotyping of the microbial pipefish gut community.

The limited presence of trans-generational plasticity gives only partial support for our hypotheses and is in strong contrast to previous experiments performed with the same model system, where the genetic background was mostly ignored and experiments focused only on one population (Roth, Keller et al. 2012, Beemelmanns and Roth 2016, Beemelmanns and Roth 2017, Roth and Landis 2017). The here performed experiment allows us to at least partially disentangle genetic adaptation and trans-generational plasticity and suggests that selection imposed by genetic adaptation is a lot stronger than the impact of trans-generational plasticity. To this end, the unexpected limited identification of trans-generational plastic effects could indicate that we are generally overestimating trans-generational plasticity in experiments that ignore genetic background, as genetic adaptation is intermingled with the phenotypic plastic components. Alternatively, we have potentially not identified all present signs of trans-generational plasticity in this experiment as the populations are too distinct due to their history of genetic adaptation hindering the identification of trans-generational plastic effects. By taking the genetic adaptation into account, we suggest that the probability to identify existing phenotypic plastic effects is lower, as the impact of phenotypic plastic effects is weaker than the impact of genetic differences among populations.

Populations that invaded a new habitat are under strong selection for genetic adaptation towards the novel environmental condition. They go through a bottleneck, which results in populations that are diverged from their ancestral populations (Johannesson, Le Moan et al. 2020) and are characterized by a reduced genetic diversity (Johannesson and Andre 2006). In another study this reduced genetic diversity as a consequence of genetic adaptation negatively impacted the individual phenotypic plasticity of sticklebacks populating low salinity regions of the Baltic Sea (DeFaveri and Merila 2014, Hasan, DeFaveri et al. 2017). In a stable salinity environment, we would thus expect that genetic adaptation had resulted in reduced phenotypic plasticity and lower performance in the ancestral environment (DeWitt, Sih et al. 1998, Schneider and Meyer 2017). In contrast to our expectation, juvenile survival of parents from low salinity origins was not reduced at high developmental salinity suggesting that genetic adaptation towards low salinity conditions did not result in a reduction of phenotypic plasticity. Along the same line, the smaller size of juveniles from parents originating from low salinity environments is no indicator for reduced plasticity either. The smaller phenotype (at the same age) was more likely a result of the reduced parental size (Nygard, Kvarnemo et al. 2019), which can be an adaptation to low salinities caused by shifts in allele frequencies (McGuigan, Nishimura et al. 2011). The strong salinity fluctuations in the coastal environments across the Baltic Sea (Bock and Lieberum 2017) most likely selected against the loss of phenotypic plasticity.

The isosmotic level of many marine fish is equivalent to around 12 PSU (Schaarschmidt, Meyer et al. 1999) or a couple of units higher, depending on the ambient salinity conditions (Quast and Howe 1980, Partridge, Shardo et al. 2007). This suggests that the here applied high salinity treatment is rather hyper-to isosmotic, whereas the low salinity treatment is hypoosmotic. The hormone prolactin is involved in many metabolic pathways in vertebrates and highly relevant for fish in hypoosmotic conditions as it prevents the loss of ions and the uptake of water. Both mechanisms are crucial in hypoosmotic conditions to maintain homeostasis (McCormick 2001, Manzon 2002, Breves, McCormick et al. 2014). In our study, prolactin (*prl*) was the gene with the strongest upregulation in juveniles at low developmental salinity conditions underlining the ability of pipefish to quickly respond to prevailing salinity conditions. Similar patterns in the upregulation of prolactin in marine fish have been identified in black porgy *Acanthopagrus schlegelii* (Tomy et al. 2009) and rainbow trout *Oncorhynchus mykiss* (Prunet et al. 1990). This implies that higher *prl* expression under low salinity conditions could be indicative for adaptive developmental plasticity and suggest that juvenile fish are able to cope with short term salinity changes.

Some strains of the species *Vibrio alginolyticus* have been shown to become more virulent under low saline conditions (Dayma, Raval et al. 2015, Poirier, Listmann et al. 2017). Drivers for this increased virulence can be trade-offs in the host (Birrer, Reusch et al. 2012, Poirier, Listmann et al. 2017), a phenotypic response of the bacteria (Hase and Barquera 2001, Dayma, Raval et al. 2015) or a genetic adaptation of the bacteria to low salinity (Brown, Cornforth et al. 2012). Under low saline condition, we thus expected strong selection for immunological adaptation towards the prevailing pathogens that potentially resulted in a higher tolerance or a more effective immune defense against *Vibrio* bacteria. In line with this expectation, we found that pipefish offspring from parents caught at low salinity origin survived better when exposed to *Vibrio* bacteria than offspring from parents caught at high saline origins. This suggests that local adaptation to low saline conditions allows pipefish to allocate sufficient resources towards their immune system for fighting *Vibrio* infections. To this end, we found support for our hypothesis that increased *Vibrio* virulence in marine host organism can result from resource allocation tradeoffs towards osmoregulation, impairing the host’s immune system (Birrer, Reusch et al. 2012).

The bacteria used in this experiment were previously evolved at the respective high (V15: 15 PSU) or low (V7: 7 PSU) Baltic Sea salinity condition. If genetic adaptation of bacteria to low salinity induces their virulence, we would have expected that the bacteria evolved at 7 PSU (V7) are more virulent, in particular for the pipefish offspring from parents originating from high saline locations. In contrast to our expectation, we have identified that v15 caused a higher mortality in juveniles originating from a high saline environment than in juveniles coming from low salinity origin and low parental salinity acclimation, while the impact of v7 was not differentiable across all groups. Gene expression measurements were not appropriate to answer the question of induced virulence and a corresponding stronger host immune response against bacteria evolved at low or high salinity depending on pipefish local salinity adaptation. Instead, the injury imposed by the injection had the strongest impact on the gene expression pattern. As such, gene expression of sham-injected animals was not distinguishable from those injected either with v7 or v15. This is an unexpected limitation of our study, however, given that impact of *Vibrio* bacteria on pipefish mortality could have been identified, we are assuming that we have not chosen the time point when reactions towards *Vibrio* infection would have been best mirrored in the gene expression patterns but rather the timepoint when inflammation or stress upon the injection could be assessed. Bearing these limitations in mind, we suggest that the increased virulence of the *V. alginolyticus* strain is mainly driven by trade-offs impairing the pipefish’s immune system. A deficiency that can potentially be overcome by local adaptation.

The here identified patterns have to be interpreted with care. Due to the unintended brood pouch infection that negatively affected 47% of the pregnant males originating from high saline conditions and parentally acclimated at low saline conditions, we are dealing with distinct selection intensities on the different treatment groups (Roth, Beemelmanns et al. 2018). In the treatment affected by the brood pouch fungus (Origin Salinity: *H*, Acclimation salinity: L) multiple clutches have at least been partially lost and potentially all infected fathers were suffering stress levels that can seriously confound the results from this study. The brood pouch fungus has severely impacted offspring development such that only the strongest will have survived. Addressing life history traits in the offspring and their gene expression will thus in the *H*L group only be done in the strongest animals, which does not resemble the original cohort, and makes interpretation of the data in the offspring generation difficult. We are aware of this limitation and have been taking this into account when interpreting our data.

## 4.6 Conclusion

After the last glacial maximum, broad-nosed pipefish have successfully populated the low salinity areas of the Baltic Sea. The results of our study suggest that the components of this success story are a mixture of genetic adaptation and the maintenance of a high degree of phenotypic plasticity of locally adapted pipefish enabling them to deal with present and ancestral salinity levels and even with re-occurring salinity fluctuations. Pipefish individuals with suitable alleles for low salinity conditions can inhabit low saline environments. The adaptation and adjustment of life history strategies to lower salinity also enable pipefish to cope with prevailing pathogens such as *Vibrio* bacteria or aquatic fungi. Pipefish of the species *S. typhle* inhabiting the Baltic Sea are thus expected to be well prepared for predicted further drops in salinity, which is also underlined by the fact that already now they inhabit areas in the northern part of the Baltic Sea with salinity levels below 5 PSU.

## Supporting information

Document with all supplement material

## Acknowledgments

We are grateful for the help of Kristina Dauven, Andreas Ebner, Janina Röckner and Paulina Urban for fish collection in the field and fish maintenance. Furthermore, we thank Fabian Wendt for setting up the aquaria system. We thank Tatjana Liese, Paulina Urban, Jakob Gismann and Thorsten Reusch for support with DNA extraction and analysis of pipefish population structure. The authors acknowledge support of Isabel Tanger, Agnes Piecyk, Jonas Müller, Grace Walls, Sebastian Albrecht, Julia Böge and Julia Stefanschitz for their support in preparing cDNA and running of Fluidigm chips. A special thank goes to Diana Gill for general lab support, ordering materials and just being the good spirit of our molecular lab, to Till Bayer for bioinformatics support and to Melanie Heckwolf for fruitful discussion and feedback on the manuscript. H.G. is very grateful for inspirational office space with ocean view provided by Lisa Hentschel and family.

## Ethical statement

Experimental work was conducted in agreement with the German animal welfare law and approved by the Ministerium für Energiewende, Landwirtschaft, Umwelt, Natur und Digitalisierung under permission MELUR V 312–7224.121-19 (67–5/13), “komparative Vergleichsstudie von Immunantwort-Transfer von Eltern zu Nachkommen in Fischarten mit extremer Brutpflege”)

## Author contribution

O.R. and H.G. designed the study with input from C.C.W. H.G., K.W., O.R. collected fish in the field, conducted the experiment and sampled the fish. L.S. designed the primers. L.S., H.G. and K.W. did the molecular lab work. H.G., L.S., O.R and K.W. analyzed the data. H.G., L.S. and O.R. wrote the manuscript with input from all others.

## Data archiving

