## Supplementary material for "Pipefish locally adapted to low salinity in the Baltic Sea retain phenotypic plasticity to cope with ancestral salinity levels": Document with all supplement material

**Supplement 1: Table S 1:** Name and function of target genes used to study gene expression. Listed are all target genes from four different functional categories: general metabolism, immune response (adaptive and innate), gene regulation (acetylation and methylation) and osmoregulation. The column 'Organism' specifies, whether the gene was targeted in female (f) or juvenile (juv) samples.

| Gene | Organism | Category | Gene name | Function | Reference |
|---|---|---|---|---|---|
| BROMO | juv, f | epigenetic/ activation | Histone acetyltransferase | Histone acetylation | Beemelmanns & Roth 2016 |
| MYST | juv | epigenetic | Histone acetyltransferase | Histone acetylation | Beemelmanns & Roth 2016 |
| Bcell.rap | juv | adaptive | B-cell receptor-associated protein | T- and B-cell regulation activity | Roth et al. 2012 |
| CD45 | juv | adaptive | CD45 (Leukocyte common antigen) | T-cell and B-cell antigen receptor signalling | Beemelmanns & Roth 2016 |
| HIVEP2 | juv | adaptive | Human immunodeficiency virus type l enhancer 2 | Transcription factor, V(D)J recombination, MHC enhancer binding | Beemelmanns & Roth 2016 |
| HIVEP3 | juv | adaptive | Human immunodeficiency virus type l enhancer 3 | Transcription factor, V(D)J recombination, MHC enhancer binding | Beemelmanns & Roth 2016 |
| IgM | juv | adaptive | Immunoglobulin light chain | Antigen/pathogen recognition | Beemelmanns & Roth 2016 |
| lymphag75 | juv | adaptive | Lymphocyte antigen 75 | Antigen recognition | Birrer et al. 2012 |
| aif | juv, f | innate | Allograft inflammation factor | Inflammatory responses, allograft rejection, macrophages activation | Roth et al. 2012 |
| apoa1 | juv | innate | Apolipoprotein A1 | Antimicrobial activity | Roth et al in prep. |
| c1 | juv | innate | Recognition subcomponent C1q | Antigen-antibody complex formation | Beemelmanns & Roth 2016 |
| c3 | juv | innate | Complement component 3 | Complement system activation | Birrer et al. 2012 |
| c9 | juv | innate | Complement component 9 | Membrane attack complex, bacteria lysis | Roth et al. 2012 |
| cf | juv, f | innate | Coagulation factor II | Blood clotting and inflammation response | Birrer et al. 2012 |
| hsp60 | juv, f | innate | Heat shock protein 60 kDa | Chaperone, general stress response | Roth et al. 2012 |
| il10 | juv | innate | Interleukin 10 | Macrophage activity regulation | Birrer et al. 2012 |
| il8 | juv | innate | Interleukin 8 | Phagocytosis, inflammation | Beemelmanns & Roth 2016 |

| Gene | Organism | Category | Gene name | Function | Reference |
|---|---|---|---|---|---|
| kin | juv, f | innate | Kinesin | Intracellular transport | Roth et al. 2012 |
| lectpt2 | juv, f | innate | Lectin protein type II | Pathogen recognition receptor | Beemelmanns & Roth 2016 |
| tranfe | juv, f | innate | Transferrin | Bacterial growth prevention | Beemelmanns & Roth 2016 |
| tspo | juv, f | innate | Translocator protein | Inflammatory responses, allograft rejection, macrophage activation | Roth et al. 2012 |
| tyroprot | juv, f | innate & adaptive | Tyroproteinkinase | Cytokine receptor signalling | Beemelmanns & Roth 2016 |
| ddpgly | juv, f | metabolism | Ddp-glycosyltransferase | Metabolizing process (natural glyosidic linkages) | Roth et al. in prep. |
| g6DPH | juv, f | metabolism | Glucose 6 phosphate dehydrogenase (G6PD) | Metabolizing process (pentose phosphate pathway) | Roth et al. in prep. |
| ptgis | juv | metabolism | Prostaglandin I2 Synthase | Lipid and fatty acid metabolism | Roth et al. in prep. |
| ripop | juv, f | metabolism | Ribosomal protein | Translation process | Roth et al. in prep. |
| TNF | juv | metabolism | Tumor necrosis factor | Lipid metabolism | Roth et al. in prep. |
| ubi | juv, f | metabolism | Ubiquitin | Regulatory protein labelling for degradation | Birrer, Reusch, et al. 2012 |
| DnMt3B | juv | epigenetic | DNA methyltransferase 3b | De novo methylation | Beemelmanns & Roth 2016 |
| JmicPhD | juv, f | epigenetic/ silencing | Lysine-specific demethylase 5B | Histone demethylation | Beemelmanns & Roth 2016 |
| N6admet | juv | epigenetic/ activation | N(6)- adenine-specific DNA-methyltransferase 2 | DNA-methyltransferase | Beemelmanns & Roth 2016 |
| no66 | juv,f | epigenetic/ silencing | Lysine-specific histone demethylase NO66 | Histone demethylation | Beemelmanns & Roth 2016 |
| TPR | juv, f | epigenetic/ activation | Lysine-specific demethylase 6A | Histone demethylation | Beemelmanns & Roth 2016 |
| aqp3 | juv | osmoregulation | Aquaporin 3 | Water and small solute channel | This study |
| atp1a1 | juv | osmoregulation | ATPase alpha 1 | Na+/K+ transporting | This study |
| cftr | juv | osmoregulation | Cystic fibrosis transmembrane conductance regulator | Apical membrane anion channel | This study |
| cldn1 | juv | osmoregulation | Claudin 1 | Epithelial permeability regulation | This study |
| hsp70 | juv | osmoregulation | Heat shock protein 70 kDa | Osmotic stress response | This study |

| Gene | Organism | Category | Gene name | Function | Reference |
|---|---|---|---|---|---|
| kcnh8 | juv | osmoregulation | Voltage gated potassium channel subfamily h member 8 | Cell volume regulation | This study |
| mapk8ip3 | juv | osmoregulation | Mitogen-activated protein kinase 8 interacting protein 3 | Osmosensing | This study |
| nkcc2 | juv | osmoregulation | Na+/K+ /2Cl cotransporter (Slc12A1) | Ion transport | This study |
| nr3c1 | juv | osmoregulation | Nuclear Receptor Subfamily 3 Group C Member 1 | Glucocorticoid receptor | This study |
| prl | juv | osmoregulation | Prolactin | Ion uptake promotion; ion secretion inhibition | This study |
| prlr | juv, f | osmoregulation | Prolactin receptor | Prolactin receptor | This study |
| hdac1 | juv | reference/ epigenetic | Histone deacetylase 1-like | Histone deacetylation | Beemelmanns & Roth 2016 |
| hdac3 | juv | reference/ epigenetic | Histone deacetylase 3-like | Histone deacetylation | Beemelmanns & Roth 2016 |
| ash | juv | reference/epigene tic/activation | Histone methyltransferase | Histone methyltransferase | Beemelmanns & Roth 2016 |
| calrcul | f | innate | Calreticulin | Chaperone, promotes phagocytosis and clearance of apoptotic cells | Beemelmanns & Roth 2016 |
| ck7 | f | innate & adaptive | Chemokine 7 | Chemotaxis for leukocytes, monocytes, neutrophils, blood cells | Beemelmanns & Roth 2016 |
| dnmt1 | f | epigenetic/ silencing | DNA (cytosine-5)-methyltransferase 1 | copies complementary marks of newly replicated DNA, maintenance methylation | Beemelmanns & Roth 2016 |
| dnmt3a | f | epigenetic/ silencing | DNA (Cytosine-5-)-Methyltransferase 3 Alpha | de novo modifications; essential for epigenetic changes based on environmental stress | Beemelmanns & Roth 2016, |
| hdac6 | f | epigenetic/ silencing | Histone deacetylase 6-like | Histone deacetylation (deacetylation lysine residues of core histones) | Beemelmanns & Roth 2016 |
| hemk2 | f | epigenetic/ silencing | HemK-methyltransferase family member 2 | DNA methyltransferase (N6-methyladenine) | Beemelmanns & Roth 2016, |
| ik-cytoine | f | adaptive & innate | Ik cytokine(RED-protein) | Inhibits interferon gamma mediated downregulation of MHCII | Beemelmanns & Roth 2016 |
| Intf | F | Innate | Interferon induced transmembrane protein 3 | Negative regulation of viral entry into host cell, antiviral response | Beemelmanns & Roth 2016 |

### Supplement 2: Description and selection of osmoregulatory genes

Osmoregulatory target genes were selected based on their function, involvement in adaptation to marine or freshwater environment, and/or inducibility upon salinity stress. The genes aquaporin 3 (*AQP3*), claudin 1 (*CLDN1*) and mitogen-activated protein kinase 8 interacting protein 3 (*MAPK8IP3*) were recently shown to alter their expression in response to different salinity treatments in alewives *Alosa pseudoharengus* (Velotta, Wegrzyn et al. 2017). *AQP3* is a member of transmembrane channel proteins transporting water and small solutes. It plays a major role in osmoregulatory organs such as the gill, kidney, oesophagus and intestine (Cutler, Martinez et al. 2007). *CLDN1* is a cell surface component of tight junction complexes (Paris, Tonutti et al. 2008) and *MAPK8IP3* is involved in osmosensing (Velotta, Wegrzyn et al. 2017). ATPase $Na^+/K^+$ transporting subunit alpha 1 (*ATP1α1*), cystic fibrosis transmembrane regulator (CFTR), $Na^+/K^+/2Cl^-$ cotransporter (*NKCC*) and heat shock protein 70 kDa (HSP70) were all found to be induced upon salinity changes (Hwang and Lee 2007, Taugbol, Arntsen et al. 2014, Ronkin, Seroussi et al. 2015). ATP1α1, a membrane spanning enzyme active for example in fish gills, actively transports sodium ions ($Na^+$) out of and potassium ions ($K^+$) into a cell (Cutler, Martinez et al. 2007, Hwang and Lee 2007). *CFTR* is an apical membrane anion channel secreting *$Cl^-$*, and *NKCC2* (NKCC isoform 2; also known as solute carrier family 12 member 1) is an ion cotransporter located in the membrane (Hwang and Lee 2007) Other important ion transporters like voltage gated potassium channel genes regulate cell volume and the subfamily h member 4 (*KCNH4*) gene was shown to be involved in freshwater adaptation in the three-spined stickleback *Gasterosteus aculeatus* (Taugbol, Arntsen et al. 2014). However, as no orthologous for *KCNH4* could be identified in the *S. typhle* genome, the closely related *KCNH8* was selected for this study instead (Gutman, Chandy et al. 2005). Additionally, *ATP1α1, CFTR, KCNH4* were proposed to be under selection in animals exposed to marine-freshwater gradients (Tomy, Chang et al. 2009, Taugbol, Arntsen et al. 2014) showed that expression of the prolactin receptor gene (*PRLR*) was correlated with adaptation to freshwater. Among other functions, the hormone prolactin (*PRL*) regulates water and electrolyte balance by promoting ion intake and inhibiting ion secretion (McCormick 2001, Manzon 2002). Gene nuclear receptor subfamily 3 group C member 1 (*NR3C1*) encodes glucocorticoid receptor, which stimulates ATPase $Na^{+}/K^{+}$ transport and proliferation and differentiation of ion transporting chloride-cells in osmoregulatory organs (Marshall, Cozzi et al. 2005). *NR3C1* expression has also been linked to salinity stress in the euryhaline fish black porgy *Acanthopagrus schlegeli* (Tomy, Chang et al. 2009) (Table S2).

### Supplement 3: Design of primers for osmoregulatory genes

Transcripts for (iv) osmoregulatory genes in *S. typhle* were identified by searching for orthologous of candidate genes in transcriptomes from other teleost fish. The transcripts were taken from the NCBI nucleotide database (https://www.ncbi.nlm.nih.gov/nuccore) and using the basic local alignment search tool BLAST (Altschul, Gish et al. 1990). The osmoregulatory candidate genes were identified in the *S. typhle* transcriptome and genome (Haase, Roth et al. 2013, Roth, Solbakken et al. 2020). Additionally, the protein product of the *S. typhle* gene transcript was verified in UniProt (The UniProt Consortium 2017). *S. typhle* specific primer pairs for 11 osmoregulatory genes were designed with Primer3web (version 4.1.0; for parameters see Table S1) and, finally, primer specificity was checked with BLAST against the *S. typhle* genome and transcriptome. Whenever possible, primers were designed to span Exon-Exon boundaries, visualized in alignment viewer and editor AliView (Larsson 2014).

The efficiency of potential primers was tested with quantitative real time polymerase chain reactions (qPCR) in a dilution series (1:10, 1:20, 1:40, 1:80, 1:160, 1:320). Primer specificity was checked again by visual evaluation of the melting curves. Only candidate primer pairs with efficiency between 79 - 106 % and standard curve slope (log quantity vs. threshold cycles) between -3.2 and -4.2 were chosen for the study (Table S3).

**Supplement table S3: Osmoregulatory gene primers.** Primer sequences and properties of all primers osmoregulatory gene primers used in this study. Hits on scaffold, hit annotation and PCR product sequence can be found in the data archive PANGEA.

| Gene | Target | Primer Sequence (forward/reverse) | PCR product length | melting temperature | efficiency [%] |
|---|---|---|---|---|---|
| aqp3 | Aquaporin 3 | CTTCCAGATCCGCAACCTACTG | 145 | 60.74 | 105.8 |
| | | GGCAAAGTTAACCGTCAAGAACA | | 59.93 | |
| atp1a1 | ATPase alpha 1 | GCTGGGAAGAGGGAAAGATGAT | 170 | 59.83 | 85.5 |
| | | GGTCGGTTCCGTACTTCCTATG | | 60.22 | |
| zcldn1 | Claudin 1 | AACGACAACACCAAAGCTTACG | 145 | 59.97 | 89.4 |
| | | ATGATCTCCACCATGACGTACG | | 59.97 | |
| mapk8ip3 | Mitogen-activated protein kinase 8 interacting protein 3 | CACGAAATCAAAGACGCCAAGT | 130 | 60.03 | 89 |
| | | GAAGCTGTTTGTTCTCCGTCAG | | 59.78 | |
| prl | Prolactin | TGGTTTTGTCCTCTTCCGCTAA | 182 | 59.89 | 88.6 |
| | | GATGTCATTGCCGCCTCTGTA | | 60.47 | |
| prlr | Prolactin receptor | CGGCTGGATCACACTCATCTAT | 188 | 59.7 | 83.9 |
| | | TTTGGGACTCAGAGCTCCATTC | | 60.03 | |
| nr3c1 | Nuclear Receptor Subfamily 3 Group C Member 1 | TCCCGTCAACAGGAAGTCTTTT | 163 | 59.83 | 73.3 |
| | | ACGTTGGCAGTGATAGAAGAGG | | 60.09 | |
| cftr | Cystic fibrosis transmembrane conductance regulator | AAGTTGGACCTCACTGACGTTT | 125 | 60.09 | 86.4 |
| | | CGCATCAAACTGGGCTTCTTC | | 60.14 | |
| kcnh8 | Voltage gated potassium channel subfamily h member 8 | CTCATCTTTGCACTCGTCAACC | 181 | 59.84 | 79.2 |
| | | GCTGGCAGTTAAACAACGACAT | | 60.03 | |
| *hsp70* | Heat shock protein 70 kDa | TGAGGGCGTTGATTTCTACACA | 124 | 59.96 | 81.6 |
| | | GCCCTTGTCCATCTTAGCATCT | | 60.16 | |
| *nkcc2* | Na+/K+ /2Cl- cotransporter (Slc12A1) | CTTCATCCTACTGGCGGCTATT | 145 | 59.96 | 98 |
| | | CGACTCTTGATTCCTGAAAGCC | | 59.32 | |

### Supplement 3: Sample size of target gene expression for females

**Supplement table S4:** Sample size of target gene expression for females, including sampling site, salinity category at origin, salinity acclimation in the lab and number of processed females

| Origin | Origin salinity | Acclimation salinity | Number of females |
|---|---|---|---|
| Flensburg | High | High | 14 |
| | High | Low | 15 |
| Falkenstein | High | High | 14 |
| | High | Low | 14 |
| Fehmarn | High | High | 12 |
| | High | Low | 13 |
| Salzhaff | Low | High | 12 |
| | Low | Low | 15 |
| Ruegen North | Low | High | 11 |
| | Low | Low | 12 |
| Ruegen South | Low | High | 12 |
| | Low | Low | 11 |

### Supplement table 4: Pairwise $F_{ST}$ of pipefish from different sampling sites

| Comparison | FST |
|---|---|
| Falck-Fehm | 0.0239 |
| Falck-Flens | 0.0162 |
| Falck-Salz | 0 |
| Falck-RuegN | 0.0064 |
| Falck-RuegS | 0 |
| Fehm-Flens | 0 |
| Fehm-RueN | 0.0102 |
| Fehm-RuegS | 0.0015 |
| Fehm-Salz | 0.0031 |
| Flens-ReugN | 0.0033 |
| Flens-RuegS | 0.0079 |
| Flens-Salz | 0 |
| RuegN-RuegS | 0 |
| RuegN-Salz | 0 |

### Supplement 5: Statistics tables and additional graphs for total length and body weight of adult pipefish

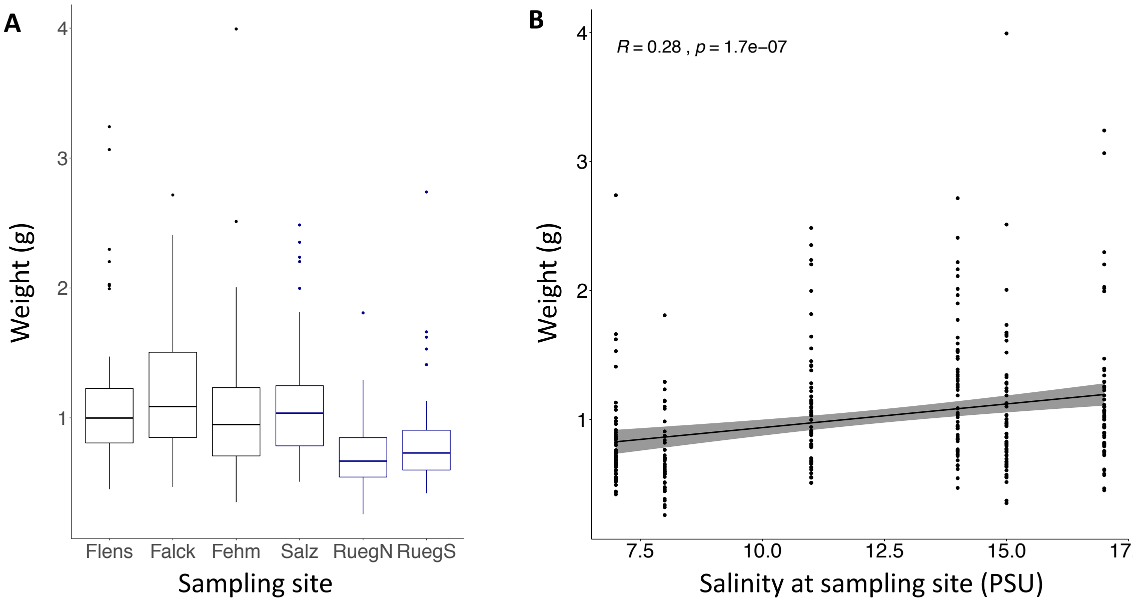


**Supplement figure S6: Weight of adult pipefish.** The weight of pipefish (y-axis) is plotted for different sampling site (x-axis) (A), and correlated with the salinity at the different sampling sites (B).

**Supplement table S6a: Analysis of variance (ANOVA) for length of adult pipefish**

Fixed factors are *Sex*, *Origin salinity*, *Acclimation salinity* and *Sampling site* nested in *Origin salinity*.

| Full model – Adult length | Df | Sum Sq | Mean Sq | F value | p |
|---|---|---|---|---|---|
| Sex | 1 | 8.82 | 8.824 | 2.4 | 0.123 |
| **Origin salinity** | 1 | 182.00 | 182.004 | 49.4 | **< 0.001*** |
| Acclimation salinity | 1 | 3.19 | 3.189 | 0.9 | 0.353 |
| Sex: Origin salinity | 1 | 1.44 | 1.443 | 0.4 | 0.531 |
| Sex: Acclimation salinity | 1 | 0.10 | 0.105 | 0.0 | 0.866 |
| **Origin salinity:Acclim salinity** | 1 | 27.27 | 27.269 | 7.4 | **0.006*** |
| **Origin salinity:Sampling site** | 4 | 166.46 | 41.616 | 11.3 | **< 0.001*** |
| Sex: Origin salinity:Acclimation salinity | 1 | 2.52 | 2.525 | 0.7 | 0.409 |
| Residuals | 320 | 1179.79 | 3.687 | | |

| Reduced model | Df | Sum Sq | Mean Sq | F value | p |
|---|---|---|---|---|---|
| **Origin salinity** | 1 | 182.0 | 182.0 | 49.4 | **< 0.001*** |
| Acclimation salinity | 1 | 3.2 | 3.2 | 0.8 | 0.353 |
| **Origin salinity:Acclimation salinity** | 1 | 26.7 | 27.3 | 7.4 | **0.007** |
| **Origin salinity:Sampling site** | 4 | 165.0 | 41.6 | 11.3 | **< 0.001*** |
| Residuals | 320 | 1179.8 | 3.7 | | |

**Supplement table S6b: Post hoc test Tukey HSD for body length of adult pipefish contrasting different salinity histories, i.e. salinity origin and acclimation salinity.**

| | **Estimate** | **Std. Error** | **t value** | **Pr(>ltl)** |
|---|---|---|---|---|
| *H*L – *H*H == 0 | -0.72 | 0.30 | -2.3 | 0.083 |
| *L*H – *H*H == 0 | -2.08 | 0.32 | -6.4 | **< 0.001*** |
| *L*L – *H*H == 0 | -1.66 | 0.32 | -5.2 | **< 0.001*** |
| *L*H – *H*L == 0 | -1.36 | 0.32 | -4.3 | **< 0.001*** |
| *L*L – *H*L == 0 | -0.94 | 0.31 | -3.0 | **0.014** |
| *L*L – *L*H == 0 | 0.41 | 0.33 | 1.2 | 0.583 |

**Supplement table S6c: Post hoc test Tukey HSD for body length of adult pipefish contrasting sampling sites.**

| | **Estimate** | **Std. Error** | **t value** | **Pr(>ltl)** |
|---|---|---|---|---|
| Fehm – Falck == 0 | -0.7373 | 0.3566 | -2.067 | 0.307 |
| Flens – Falck == 0 | -0.1271 | 0.3537 | -0.359 | 0.999 |
| **RuegN – Falck == 0** | -2.8133 | 0.3765 | -7.471 | **< 0.001*** |
| **RuegS – Falck == 0** | -2.1409 | 0.3684 | -5.811 | **< 0.001*** |
| Salz – Falck == 0 | -0.4729 | 0.3666 | -1.290 | 0.790 |
| Flens – Fehm == 0 | 0.6102 | 0.3537 | 1.725 | 0.516 |
| **RuegN – Fehm == 0** | -2.0760 | 0.3765 | -5.513 | **< 0.001*** |
| **RuegS – Fehm == 0** | -1.4036 | 0.3684 | -3.810 | **0.002*** |
| Salz – Fehm == 0 | 0.2644 | 0.3666 | 0.721 | 0.979 |
| **RuegN – Flens == 0** | -2.6862 | 0.3738 | -7.187 | **< 0.001*** |
| **RuegS – Flens == 0** | -2.0138 | 0.3656 | -5.508 | **< 0.001*** |
| Salz – Flens == 0 | -0.3459 | 0.3637 | -0.951 | 0.933 |
| ReugS – RuegN == 0 | 0.6724 | 0.3877 | 1.734 | 0.510 |
| **Salz – RuegN == 0** | 2.3404 | 0.3860 | 6.063 | **< 0.001*** |
| **Salz – RuegS == 0** | 1.6680 | 0.3781 | 4.412 | **< 0.001*** |

### Supplement 6: Statistic tables and additional graphs for clutch size

**Supplement table S7a: Analysis of variance (ANOVA) for clutch size of adult pipefish**

Fixed factors are *Origin salinity*, *Acclimation salinity* and *Sampling site* nested in *Origin salinity*.

| Full model | **Df** | **Sum Sq** | **Mean Sq** | **F value** | **p** |
|---|---|---|---|---|---|
| Origin salinity | 1 | 5.0 | 5.0 | 2.0 | 0.160 |
| **Acclimation salinity** | 1 | 41.2 | 41.2 | 15.0 | **0.001*** |
| **Av male length** | 1 | 14.6 | 14.6 | 5.8 | **0.017*** |
| **Origin salinity:Acclimation salinity** | 1 | 25.6 | 25.6 | 9.0 | **0.003*** |
| Origin salinity:Sampling site | 2 | 9.1 | 4.6 | 1.8 | 0.168 |
| Origin salinity:Av male length | 1 | 0.4 | 0.4 | 0.1 | 0.705 |
| Acclimation salinity:Av male length | 1 | 1.3 | 1.3 | 0.5 | 0.474 |
| Origin sal: Acclim sal:Av male length | 1 | 31.7 | 31.7 | 12.6 | **0.001*** |
| Residuals | 109 | 273.6 | 2.5 | | |

**Supplement table 7b: Post hoc test Tukey HSD for clutch size of adult pipefish salinity histories.** The italic letter represents the origin salinity (*H* - High, *L* - Low) and the second letter indicates the salinity level in the lab (H - High, L - Low).

| | **Estmate Std.** | **Error** | **t value** | **p** |
|---|---|---|---|---|
| ***H*L - *H*H == 0** | -22.4 | 4.6 | -4.8 | **< 0.001*** |
| ***L*H - *H*H == 0** | -14.0 | 4.5 | -3.1 | **0.010*** |
| ***L*L - *H*H == 0** | -16.5 | 4.5 | -3.6 | **0.003*** |
| *L*H - *H*L == 0 | 8.4 | 4.8 | 1.7 | 0.512 |
| *L*L - *H*L == 0 | 5.8 | 4.8 | 1.2 | 0.762 |
| *L*L - *L*H == 0 | -2.5 | 4.7 | -0.5 | 0.976 |

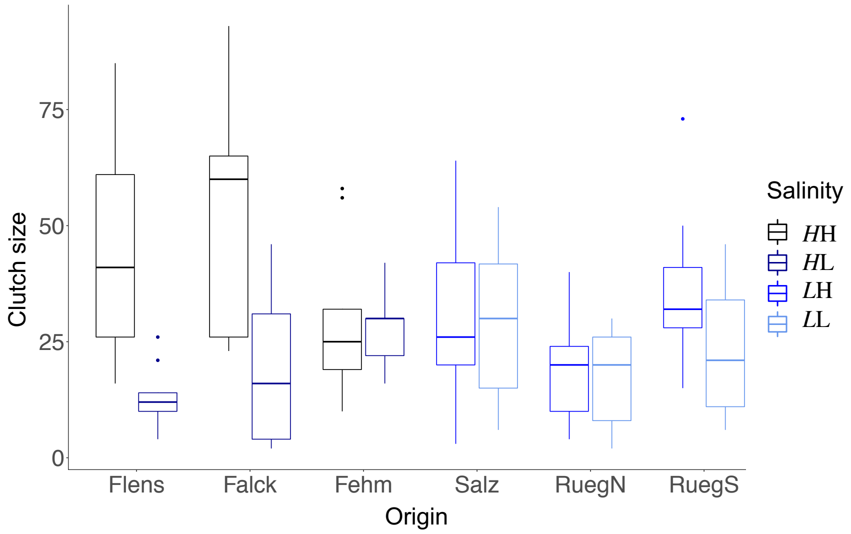


**Supplement Figure 7c: Clutch size is reduced at low salinity for all sampling sites**
Number of juveniles (clutch size) is shown for pipefish from different origins (sampling site). Italic letters and colors indicate the salinity level at the origin of (*H*: 15 PSU, black & dark blue; *L*: 7 PSU blue & light blue). The 2nd letter indicates the salinity during breeding in the lab (H: 15 PSU, black and blue; L: 7 PSU, dark & light blue).
